# Sun4 is a type II transmembrane protein of the spermatid inner nuclear membrane that forms heteromeric assemblies with Sun3 and interacts with Lamin B3

**DOI:** 10.1101/2022.04.13.488151

**Authors:** Hanna Thoma, Luisa Grünewald, Elisabeth Pasch, Manfred Alsheimer

## Abstract

SUN domain proteins are conserved proteins of the nuclear envelope and key components of the LINC complexes (linkers of the <u>n</u>ucleoskeleton and the <u>c</u>ytoskeleton). Previous studies have demonstrated that the testis-specific SUN domain protein Sun4 is a vital player in spermatogenesis, critically involved in the directed shaping of the spermatid nucleus. Its molecular properties relating to this crucial function, however, have remained largely unknown. Previous studies presented quite controversial data for the general organization and orientation of Sun4 within the spermatid nuclear envelope. In the present study, we have re-evaluated this issue in detail and present new robust data on the Sun4 topology and its interactions at the nucleo-cytoplasmic junction. We identified Sun4 as an integral protein of the inner nuclear membrane, sharing a classical SUN domain protein topology. Similar to other SUN domain proteins, the C-terminal SUN domain of Sun4 locates to the perinuclear space and the N-terminus is directed to the nucleoplasm, where it interacts with the spermiogenesis specific Lamin B3. We found that Sun4 in its natural environment forms heteromeric assemblies with Sun3 and, beyond this, it is crucially involved in the regulation of Sun3 expression. Together, our results contribute to a better understanding of the specific function of Sun4 at the spermatid nucleo-cytoplasmic junction and the entire process of sperm-head formation.

**Summary statement:** In our current study, we have analyzed in detail the biochemical and dynamic properties of the testis-specific SUN domain protein Sun4 and we provide novel insights into its interaction behavior at the spermatid nucleo-cytoplasmic junction.

## Introduction

During differentiation, most eukaryotic cells undergo fundamental changes in their function and morphology. Besides phenotypical changes of the cell itself, cellular differentiation often involves an (active) relocation and restructuring of the nucleus (Burke and Roux, 2009; Fridkin et al., 2009; Skinner and Johnson, 2017; Starr, 2009). One example of very pronounced differentiation-related nuclear restructuring occurs during spermiogenesis, which describes the maturation of haploid spermatids into fertilization-competent spermatozoa. Apart from the formation of the flagellum and the acrosome, spermiogenesis is primarily characterized by a prominent nuclear reshaping from round to elongated (Hermo et al., 2010; Hermo et al., 2009; Russell et al., 1991; Kierszenbaum and Tres, 2004; Kierszenbaum et al., 2003). Nuclear elongation, in turn, is a highly organized process combining different cellular processes, including the assembly of sperm(atid)-specific cytoskeletal structures, nuclear movement, compaction of the chromatin, and extensive remodeling of the nuclear envelope (NE), which comprises alterations of the general NE composition, as well as relocalization of various NE components. Best examples are the Lamins B1 and B3, the lamina associated protein Lap2, and the Lamin B receptor, which characteristically polarize to the posterior pole of the spermatid nucleus with the progression of spermiogenesis (Alsheimer et al., 1998; Mylonis et al., 2004; Schütz et al., 2005a; Vester et al., 1993). Several recent studies suggest that LINC (linkers of the <u>n</u>ucleoskeleton and the <u>c</u>ytoskeleton) complexes have a very central function in spermatid nuclear remodeling and elongation (Calvi et al., 2015; Gao et al., 2020; Göb et al., 2010; Pasch et al., 2015; Yang et al., 2018). LINC complexes are conserved NE-bridging assemblies that physically connect the nucleus to the peripheral cytoskeleton, thus providing a molecular basis for the transfer of mechanical forces to the NE and beyond (Crisp et al., 2006; Starr and Fridolfsson, 2010). The central core of the LINC complexes is formed by interaction of members of two transmembrane protein families: The SUN (Sad-1/UNC-84) domain proteins and the KASH (Klarsicht/ANC-1/Syne/homology) domain proteins (Crisp et al., 2006; Razafsky and Hodzic, 2009; Starr and Fridolfsson, 2010; Stewart-Hutchinson et al., 2008). SUN domain proteins are highly conserved proteins of the inner nuclear membrane (INM) which share a common C-terminal SUN motif (Hagan and Yanagida, 1995; Malone et al., 1999). They are known to N-terminally anchor to structural components of the nuclear interior, e.g., the nuclear lamins, while their C-terminal part which mainly consists of a central coiled-coil motif and the conserved SUN domain, locates within the perinuclear space (PNS) (Hodzic et al., 2004; Padmakumar et al., 2005). In mammals, five different SUN proteins are known: Sun1 and Sun2, which are expressed in all kinds of somatic cells (Hodzic et al., 2004; Padmakumar et al., 2005), and Sun3, Sun4 (Sapg4) and Sun5 (Spag4L), which are exclusively expressed in spermatids (Calvi et al., 2015; Frohnert et al., 2011; Göb et al., 2010; Pasch et al., 2015). KASH domain proteins (Nesprins), for their part, are integral to the outer nuclear membrane (ONM) with their large N-terminal domains (NTDs) connected to the cytoskeleton and their short C-terminal KASH domain protruding into the PNS, where they specifically interact with the SUN domain of their SUN protein binding partner (Sosa et al., 2012; Starr, 2009). In their role as connectors of nuclear and cytoskeletal structures, LINC complexes are key players in various developmental cellular processes involving nuclear movement, positioning or anchorage, cell shaping, the establishment of cell polarity, cell migration, meiotic chromosome movement, nuclear shaping and fertilization (Kracklauer et al., 2013; Razafsky and Hodzic, 2009; Starr and Fridolfsson, 2010).

Within the last decade, evidence has been accumulating that nuclear shaping and the formation of a functional sperm-head during spermiogenesis vitally depends on the cooperative function of a variety of different LINC complex components. According to their designated functions, these components selectively localize to distinct territories of the differentiating nucleus. Sun3, for example, interacts with Nesprin1, which both locate to the posterior NE in round and elongated spermatids. Sun3/Nesprin1 are found exclusively in regions where the manchette microtubules are in close contact with the NE (Göb et al., 2010; Gao et al., 2020). A similar NE localization has been described for Sun4, which colocalizes with Sun3 at the posterior NE during the entire differentiation process (Pasch et al., 2015, Calvi et al., 2015). The distribution of Sun5, however, still remains rather ambiguous. It initially was suggested to be part of the anterior NE, where it was believed to be involved in acrosome biogenesis (Frohnert et al., 2011), while in subsequent studies, it was identified as a part of the posterior NE and the head-to-tail coupling apparatus (HTCA), where it is most likely involved in regulating head-to-tail linkage (Shang et al., 2017; Yassine et al., 2015; Zhang et al., 2021). In addition to the testis-specific SUN domain proteins the rather ubiquitously expressed Sun1 and Nesprin3 are also present in the early spermatid stages. They both appear to interact with each other and transiently localize to the very posterior pole, from which they gradually disappear with advancing differentiation. Concomitantly, Sun1η, a short isoform of the Sun1 gene, appears together with Nesprin3 and localizes to the anterior nuclear region, most likely being part of the acrosomal membrane (Göb et al., 2010).

Previous Sun4 knock-out studies demonstrated that Sun4 is essential for correct positioning of other NE components, proper manchette assembly, and thus, also for nuclear elongation and accurate sperm-head formation (Calvi et al., 2015; Pasch et al., 2015; Yang et al., 2018). A similar function was recently also described for Sun3. The Sun3 knock-out phenotype perfectly mirrors that of Sun4, indicating that both are involved in the same process. This is supported by the fact that Sun3 and Sun4 depends on each other for their correct localization (Calvi et al., 2015; Pasch et al., 2015; Gao et al., 2020). Based on this mutual dependency, it seems very likely that Sun3 and Sun4 cooperate in forming functional LINC complexes with Nesprin1 to physically link the manchette microtubules to the NE.

In the case of Sun4, the particular molecular properties relating to its function are still largely unclear. According to its basic composition, Sun4 overtly resembles a typical SUN domain protein. It is expected to be an integral protein of the INM, N-terminally connecting to the nucleoskeleton, whereby its C-terminal part is directed to the PNS (Meinke and Schirmer, 2015). However, some previous studies suggested Sun4 to be also located at the axoneme and the HTCA. At these sites, based on evidence from *in vitro* experiments, Sun4 was assumed to bind via its SUN domain to the outer dense fiber protein Odf1 (Shao et al., 1999; Yang et al., 2012; Yang et al., 2018). This scenario is in clear conflict with the generally accepted *bona fide* membrane topology of SUN domain proteins, as the implied typical PNS localization of the SUN domain would make direct interactions with cytoskeletal elements basically impossible. To date, it has *de facto* never been validated how Sun4, in reality, is oriented within the NE; its actual localization within the NE and its intrinsic membrane topology has remained rather hypothetical. Beyond this, it is also still not clear how Sun4 is organized within a functional LINC complex and how this complex as a whole might anchor to the spermatid nucleoskeleton.

To better understand the actual function and basic structure of Sun4 containing LINC complexes, we conducted a detailed analysis of Sun4 with the aim of defining its factual membrane topology and *in vivo* interaction behavior. Consistent with all as yet analyzed SUN domain proteins, we here characterize Sun4 as a typical type II membrane protein that is integral to the INM, with its C-terminal SUN domain located within the PNS and its N-terminal part reaching into the nuclear interior of the spermatid nucleus. In addition, we provide clear evidence that of the two apparent hydrophobic motifs that were identified in the Sun4 amino acid sequence, only one is a functional transmembrane domain (TMD), while the other one is not integral but confers membrane affinity and adds to the overall membrane retention of the protein. Furthermore, we show that Sun4 forms heteromeric assemblies with Sun3 in its natural context, i.e. within spermatids, and that it anchors to the nuclear lamina via a specific interaction of its NTD with Lamin B3. Thus, our results provide further important insights into the specific function and behavior of Sun4 and its role in the formation of functional LINC complexes.

## Results

### Sun4 is a type II transmembrane protein

Common transmembrane prediction tools (e.g., TMHMM or SOSUI; http://www.expasy.org/proteomics) indicate the presence of two hydrophobic, putative membrane-spanning domains within the N-terminal half of the Sun4 protein sequence of both, mouse (HM1: amino acids 137-159; HM2:166-188) and man (HM1: 135-157; HM2: 167-189), respectively (Fig. 1). It is, however, unclear whether Sun4 is a true transmembrane protein at all. Furthermore, it has not yet been experimentally determined which of the two domains is factually membrane-spanning and how Sun4 is *de facto* oriented within the NE.

**Fig 1.**
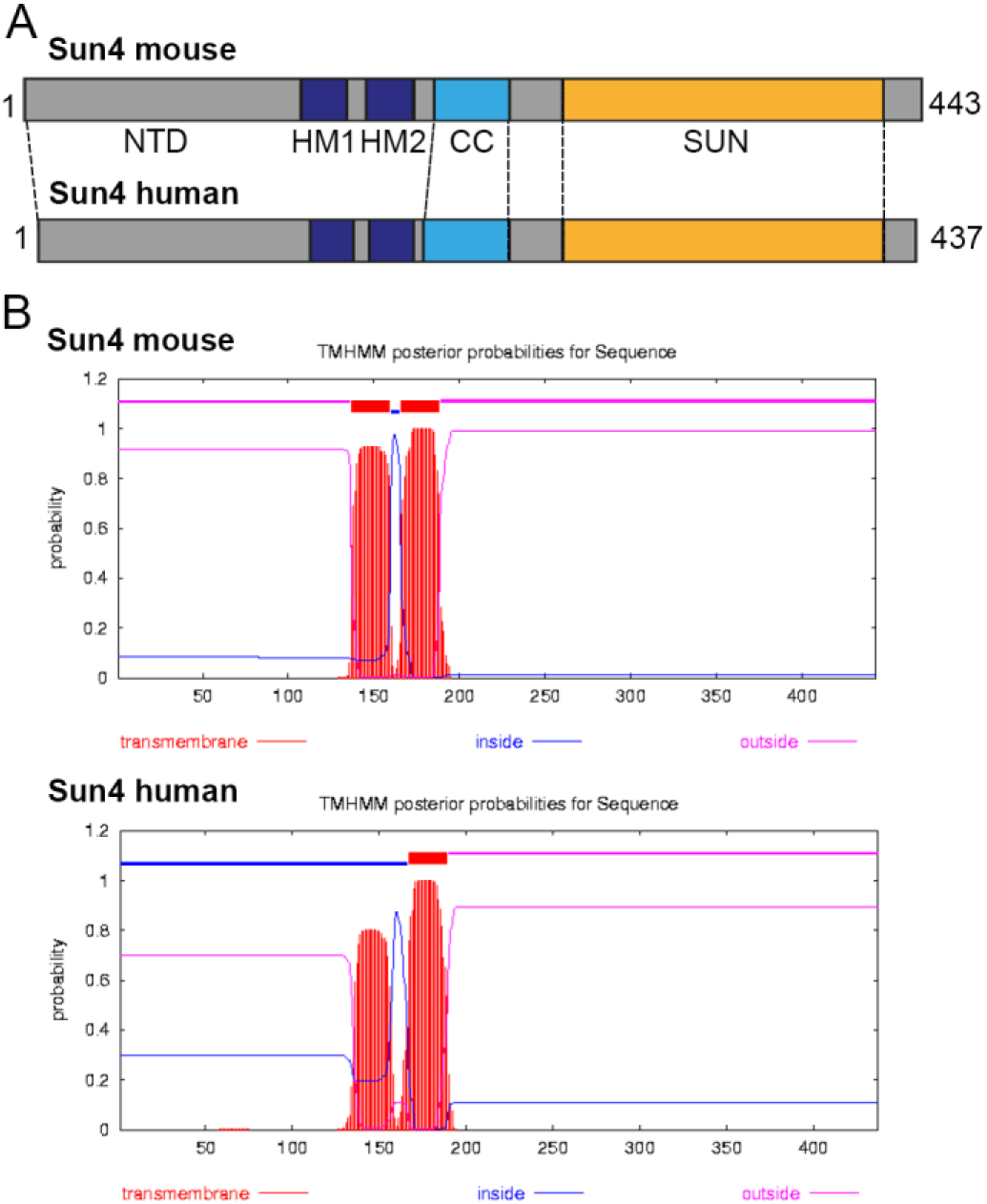
Sun4 features two hydrophobic, potentially membrane-spanning elements. (A) Schematic illustration of human (h) and mouse (m) full-length Sun4 proteins. (B) TMHMM prediction tool (http://www.cbs.dtu.dk/services/TMHMM/) identifies two hydrophobic, putative transmembrane elements, HM1 and HM2. HM1 (h: amino acids (aa) 135-157; m: aa 137-159); HM2 (h: aa 167-189; m: aa 166-188). NTD, N-terminal domain; CC, coiled-coil domain; SUN, SUN domain.

To address this issue, we started with a systematic biochemical screen largely following the protocols that were successfully applied to define the transmembrane domains (TMDs) of Sun1 and Sun5 (Frohnert et al., 2011; Liu et al., 2007). Analogous to the previous strategies, we generated a set of different Sun4 constructs – each as Myc/EGFP-tagged and untagged versions – coding for either the wild-type or mutant proteins with deletions of the C-terminal domain (CTD) and either one of the putative TMDs, HM1 or HM2 or both (Fig. 2).

**Fig 2.**
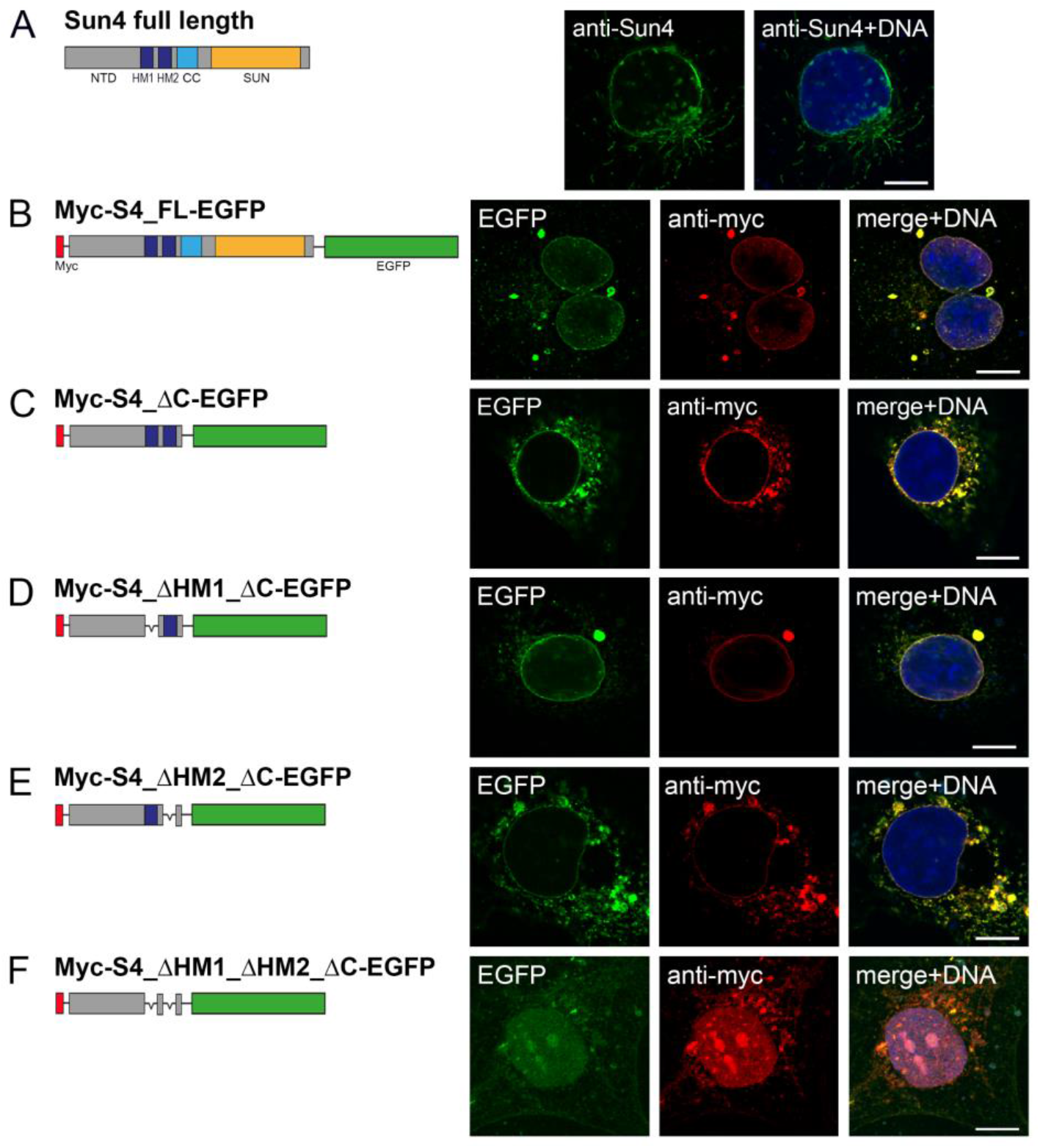
HM1 and HM2 both confer membrane affinity to Sun4. Wild-type mouse Sun4 and related Myc-/EGFP-tagged fusion constructs were transiently transfected into COS-7 cells and their respective expression patterns were analyzed 24 hours after transfection. Confocal images of (A) Sun4 and (B-F) Myc immunostainings are shown in combination with the EGFP-fluorescence. DNA was counterstained with Hoechst. Scale bars: 10 µm.

First, we transfected COS-7 cells with untagged wild-type or mutant Sun4 constructs and analyzed the behavior of the ectopically expressed proteins by immunofluorescence microscopy 24 hours after transfection. Consistent with what may be expected for a SUN domain protein, wild-type Sun4 displayed typical membrane affinity and localized to the NE and, likely due to ectopic overexpression, to the cytoplasmic membrane systems (Fig. 2A). Virtually the same behavior was observed with a Sun4 version flanked by an N-terminal Myc-tag and a C-terminal EGFP-tag (Myc-S4_FL-EGFP), suggesting that the tags do not interfere with membrane association *per se* (Fig. 2B). Likewise, a truncated Sun4 construct containing both putative TMDs but lacking the entire C-terminal, presumably perinuclear domain (Myc-S4_ΔC-EGFP), was still found to localize to the endogenous membrane systems showing a clear tendency of accumulation at the NE (Fig. 2C). Notably, single deletions of either of the two hydrophobic domains (Myc-S4_ΔHM1_ΔC-EGFP, Myc-S4_ΔHM2_ΔC-EGFP) did not prevent membrane association of the respective Sun4 constructs, indicating that each of the hydrophobic motifs, to some extent, confers membrane affinity. However, deletion of both hydrophobic motifs completely impeded membrane association, thus demonstrating that at least one of the two putative TMDs must be present to allow efficient targeting Sun4 to membranes (Fig. 2D-F).

To narrow down the factual membrane-spanning domain(s) in Sun4 and to define its effective membrane topology, we then expressed selected Sun4 constructs and performed *in situ* proteinase K digestion assays according to established protocols (Frohnert et al., 2011; Liu et al., 2007). In brief, COS-7 cells were transfected with either of the respective Myc/EGFP-tagged constructs (Figs 2, 3). 24 hours after transfection, the cells were prepared for proteinase K digestion as follows: One set of each transfection batch was directly subjected to proteinase K digestion. A second set of cells was permeabilized with 24 µM digitonin (10 minutes on ice) and subsequently incubated with proteinase K, and a third set was treated with proteinase K in the presence of 0.5% Triton-X100. The 0.5% Triton-X100 used in this assay permeabilizes the cells’ entire membrane system, whereas treatment with 24 µM digitonin selectively permeabilizes the plasma membrane while keeping the endogenous membrane systems intact. Thus, when treating the cells with 24 µM digitonin, protein parts that are located inside the endogenous membrane systems (such as the ER and the perinuclear space) are expected to be protected from proteinase K, whereas in the presence of Triton-X100, the protease has access to all parts of the protein and therefore should be able to digest membrane proteins *in extenso* (Liu et al., 2007; Frohnert et al., 2011). Accordingly, we found ectopically expressed EGFP and the INM associated nuclear lamins that we used as a control for cytoplasmic or peripheral membrane-associated proteins, entirely degraded after permeabilization with either 0.5% Triton X-100 or 24 µM digitonin. The ER-luminal disulfide-isomerase (PDI), however, was degraded when using 0.5% Triton X-100 for permeabilization but was protected from protease digestion when using 24 µM digitonin (Fig. 3C).

**Fig 3.**
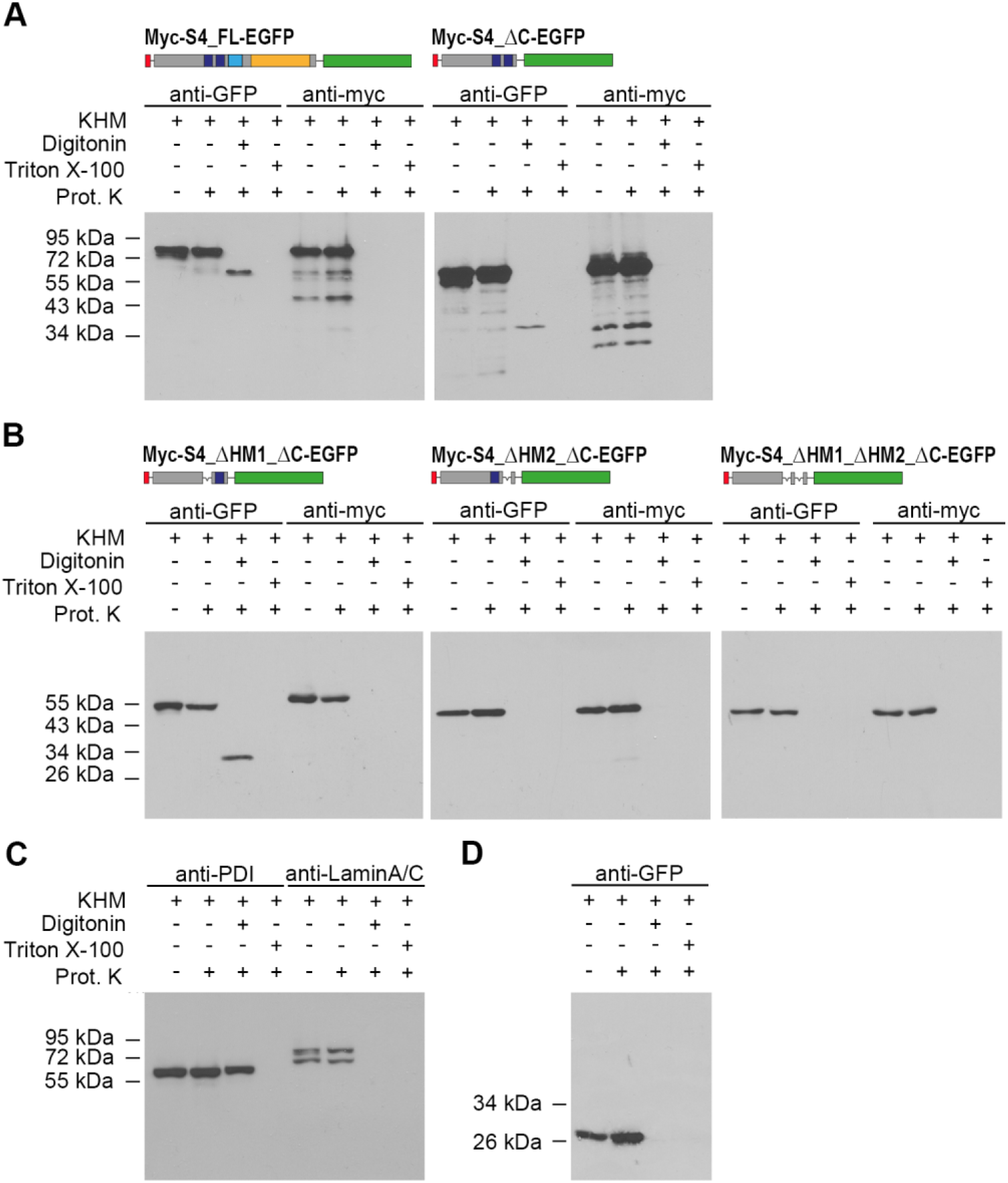
Identification of real TM domain(s) by *in situ* proteinase K digestion assay. Cos-7 cells were transfected with Myc/EGFP-tagged Sun4 constructs with deletion of (A) the C-terminal coiled-coil (CC) and (B) either HM1, HM2 or both. The cells were subjected to proteinase K digestion under different membrane permeabilization conditions and samples were processed for immunoblot analysis (lanes 2-4; first lanes show controls with untreated cells) with anti-Myc and anti-GFP antibodies. (C) Detection of PDI and Lamin A/C in untransfected cells, and (D) of EGFP in pEGFP-transfected cells, subjected to the same experimental permeabilization procedures.

Using this assay, we first tested full-length Sun4 (Fig. 3A). Here, anti-Myc and anti-GFP antibodies both detected the undigested Myc/EGFP-tagged full-length protein (apparent molecular weight of approx. 80 kDa) in untreated cells as well as in proteinase K exposed cells having intact membrane systems. In contrast, in transfected cells pre-permeabilized with 24 µM digitonin, the N-terminal Myc-tag signal disappeared entirely after proteinase K treatment, demonstrating that with permeabilization of only the plasma membrane, the NTD became accessible for protease digestion. Under the same experimental conditions, the anti-GFP antibody still recognized a peptide that appeared significantly smaller (apparent MW approx. 60 kDa), probably representing a shortened fragment lacking the NTD (approx. 18 kDa). Thus, after treatment with 24 µM digitonin, the EGFP-epitope, seemed to be protected from protease digestion, suggesting that the C-terminus of the Sun4 protein is located within the lumen of the endogenous membrane systems (Fig. 3A). As expected, proteinase K treatment after permeabilization with Triton X-100 led to degradation of the entire protein. Parallel experiments with Myc/EGFP-tagged Sun4 lacking the coiled-coil region and the SUN domain yielded virtually the same results as obtained with the Myc/EGFP-tagged full-length version, demonstrating that the C-terminal part of Sun4 has no impact on the general Sun4 topology (Fig. 3A). Together, the results presented here provide clear evidence that Sun4 is a true type II transmembrane protein. Accordingly, its membrane topology is comparable to that found in other known SUN domain proteins, with the C-terminus located within the PNS and the N-terminus protruding to the nuclear interior or the cytoplasm (Meinke and Schirmer, 2015).

### HM2 is the only valid transmembrane domain of Sun4

As a type II transmembrane protein, Sun4 requires at least one functional TMD. As mentioned above, mouse and human Sun4 protein sequences contain two hydrophobic motifs (HM1 and HM2), which, according to transmembrane prediction tools, both have a high probability of being membrane-spanning (Fig. 1). However, according to our above findings, only one can factually be a functional TMD (otherwise, the N- and C-termini would have to be situated in the same compartment). To test which of these motifs is a valid TMD, we next performed proteinase K digestion assays on cells expressing mutant Sun4 proteins that either lack HM1 or HM2 or both.

As shown in Fig. 3 B, the construct lacking both hydrophobic motifs is, as expected, completely degraded by the protease after digitonin treatment, demonstrating that at least one of the hydrophobic motifs is required for membrane integration (see also Fig. 2). In the case of the Sun4 construct lacking HM1 but still containing HM2, the EGFP-tagged C-terminus is protected from protease digestion after digitonin permeabilization. Under the same experimental conditions, the protein with intact HM1 but missing HM2, however, was completely digested. This result clearly demonstrates that of the two predicted putative TMDs, only HM2 is functional, whereas HM1 confers membrane affinity, as already indicated by the immunofluorescence experiment (Fig. 2E), but is not integral. Thus, the present experiment further substantiates that Sun4, like the other SUN protein family members, shows a typical type II membrane protein conformation consisting of a single TMD, the C-terminus located to the PNS and the NTD oriented to the nuclear interior or the cytoplasm (Meinke and Schirmer, 2015).

### The N-terminal domain of Sun4 is located within the nucleoplasm

Previous studies described that besides its localization to the posterior NE, Sun4, to some extent, also resides at the axoneme, where its CTD is presumed to interact with the outer dense fiber protein Odf1 (Shao et al., 1999; Yang et al., 2012). This scenario, however, is in clear conflict with our above finding that the C-terminal part of Sun4 is protected from protease digestion in cells with intact ER/PNS membrane systems and, thus, locates to the ER/PNS lumen. This protein part should therefore not be able to directly interact with cytoskeletal proteins such as Odf1.

In order to verify the actual localization and topology of Sun4 in its natural context, i.e. the haploid spermatids, we performed immunogold EM analysis on mouse testis tissue sections, using primary antibodies directed against the Sun4 NTD. In elongating spermatids, we found gold particles exclusively at the nucleoplasmic surface of the INM. (Fig. 4). Consistent with our earlier immunofluorescence data (Pasch et al., 2015), the gold particles decorated the central and posterior parts of the NE, which correspond to regions covered by the microtubule (MT) manchette (Fig. 4A). In late elongating spermatids, we also found considerable amounts of gold particles within the posterior, chromatin-less membrane protrusions of the redundant nuclear envelope (RNE) (Fawcett and Ito, 1965; Ho, 2010; Kerr, 1991) (Figs 4B, S1). Notably, despite using antibodies from different species and applying them on testes tissue samples fixed with different protocols, we could not detect any appreciable gold accumulations in the axoneme, as was reported in the previous immunogold study by Shao et al. (Shao et al., 1999). Our approaches demonstrate that Sun4 is solely a nuclear envelope protein of the INM. This distribution is consistent with our previous finding that Sun4 exclusively locates to the posterior NE but is excluded from the fossa region as well as the flagellum (Pasch et al., 2015; see also Fig. 7A).

**Fig 4.**
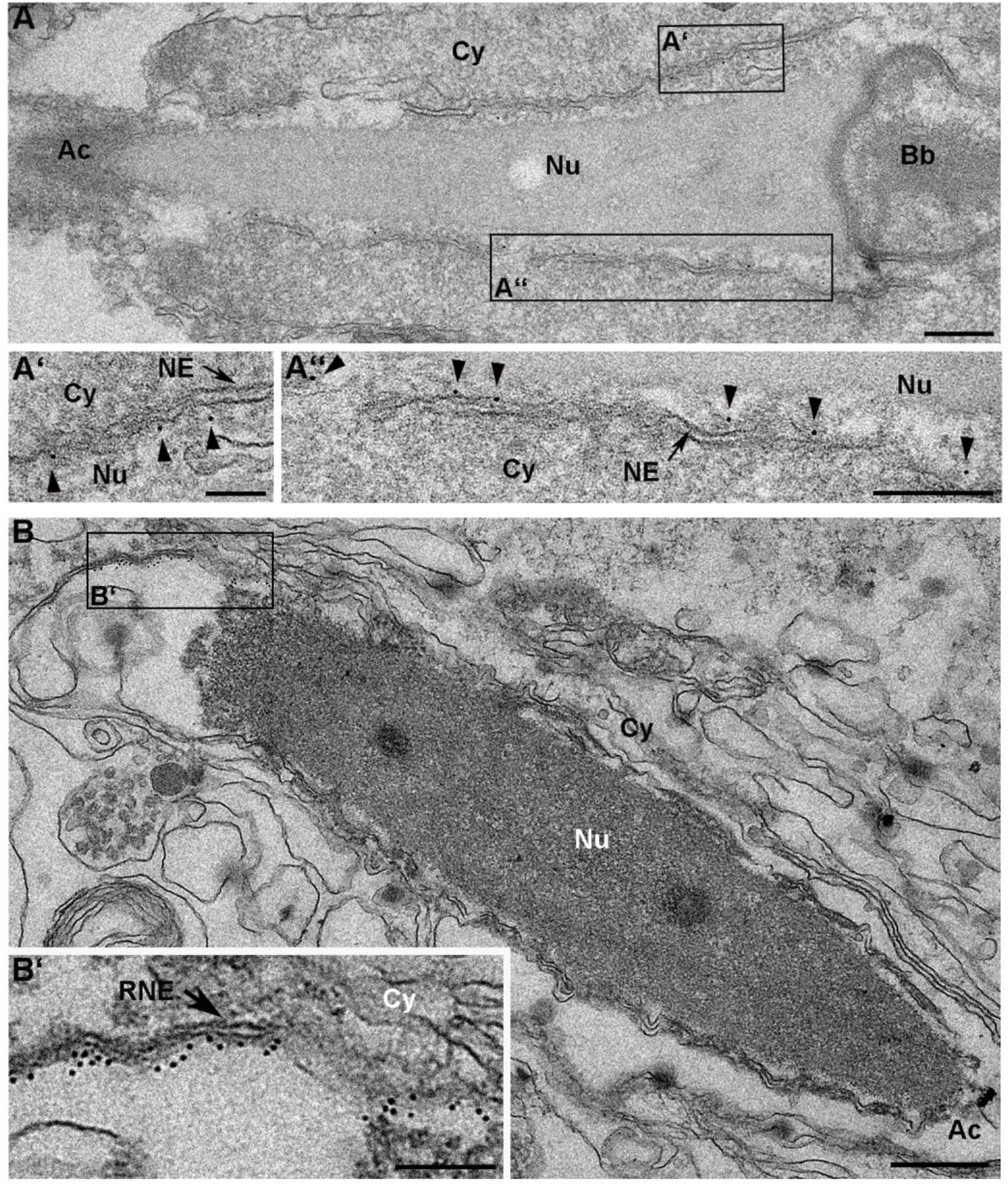
The Sun4 N-terminal domain is located within the nucleoplasm of elongating spermatids. Representative electron micrographs of elongating spermatids in (A) pre-fixed and (B) native mouse testis cryo-sections with immunogold-labelling (6 nm colloidal gold) of the Sun4 NTD. Gold particles (arrowheads) are mainly found along the inner surface of the INM (A) and the RNE (B). Higher magnifications of the areas indicated by black rectangles are depicted in A’, A” and B’. Scale bars: (B) 500 nm; (A, A”, B’) 200 nm; (A’) 100 nm. Cy, cytoplasm; Nu, nucleoplasm; NE, nuclear envelope; RNE, redundant nuclear envelope; Ac, acrosome; Bb, basal body.

In summary, our observations again confirm that the NTD of Sun4 is oriented to the nuclear interior and that Sun4, also in its native environment, shares a SUN domain protein typical type II transmembrane protein conformation (Meinke and Schirmer, 2015).

### Sun4 is exceptionally mobile within the NE of somatic cells

As we could demonstrate that Sun4, like the other SUN proteins, is integral to the INM, we next tested whether it shows comparable dynamic behavior within the membrane. To address this issue, we ectopically expressed EGFP-tagged full-length Sun4 (S4_FL-EGFP) in NIH 3T3 cells (see Fig. S2) and used FRAP to analyze its relative mobility compared to EGFP-tagged full-length Sun1 (S1_FL-EGFP) (Fig. 5B, C). Consistent with previous data published by Liu et al. (2007) and Hasan et al. (2006), Sun1-EGFP was relatively immobile and showed a low recovery of only 20.5 ± 3.9% (mean ± s. d.) of the initial pre-bleach intensity at the end of the measurement, i.e. 270 s post bleach. With the same experimental setup, S4_FL-EGFP showed much higher mobility compared to S1_FL-EGFP. As early as 10 s post-bleaching, it recovered to approximately 30% of its pre-bleach intensity, and its mean recovery rate 270 s post-bleaching even reached 90.5 ± 6.9% (Fig. 5B). Such high recovery dynamics clearly indicate that Sun4, although mostly still effectively retained within the INM (Fig. S2; see also Fig. 2B), has a low rate of local retention, at least in a heterologous, somatic environment.

**Fig 5.**
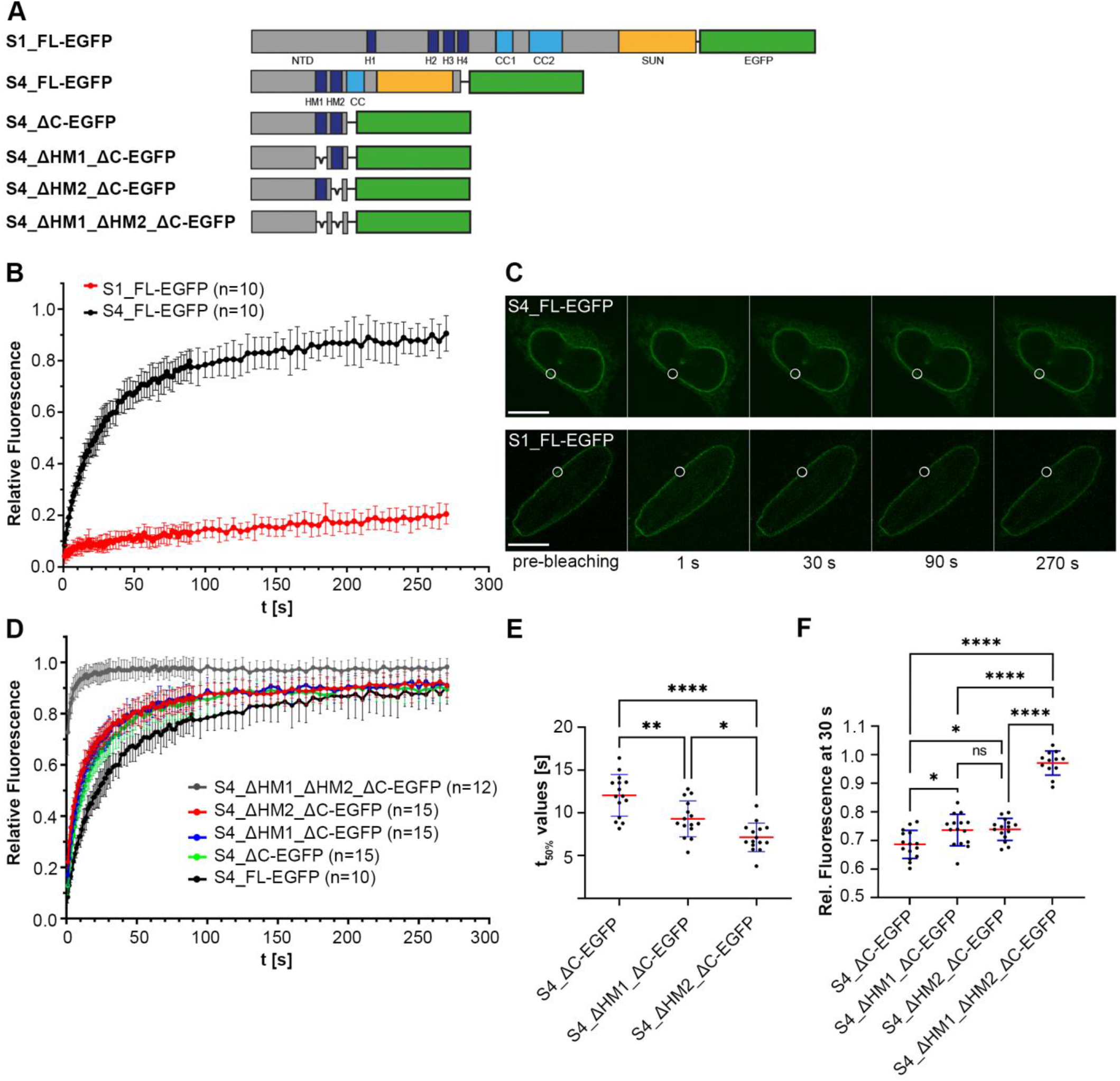
Dynamic behavior of full-length Sun4 and selected deletion mutants expressed in NIH 3T3 cells. (A) Schematic illustration of EGFP-tagged full-length (FL) Sun1, Sun4 and Sun4 deletion constructs used for FRAP analysis. (B) Recovery kinetics of S1_FL-EGFP and S4_FL-EGFP. Presented values are normalized means ± standard deviation (s.d.) of ten representative cells for each construct. (C) Time series of representative S1_FL-EGFP and S4_FL-EGFP expressing cells during FRAP-analysis. Bleaching ROIs are indicated by white circles. Scale bars: 10 µm. (D) Recovery kinetics of EGFP-tagged Sun4 deletion mutants. Presented values are normalized means ± s.d. of 10-15 (= n) representative cells per construct. Corresponding times of 50% signal recovery (t_50%_-values) and relative signal intensities 30 s post-bleaching are presented in (E) and (F), respectively. Blue error bars symbolize s.d.; red middle lines represent means. Statistical significance was tested with one-way ANOVA followed by Tukey’s multiple comparisons test. ns, not significant (p>0.05); *p≤0.05; **p≤0.01; ***p≤0.001; ****p≤0.0001.

### HM1 confers membrane association and affects the Sun4 dynamic behavior

As shown and described above, HM1, although not a factual TMD, still confers membrane affinity and, when representing the only hydrophobic motif present, efficiently targets Sun4 to the endogenous membrane system (Fig. 2E). Therefore, we next aimed to identify the particular impact of HM1 on the overall membrane behavior of Sun4. In this regard, we generated another set of Sun4 deletion constructs containing either one, both, or none of the hydrophobic elements (see Figs 5A, S2) and studied their dynamic properties within the NE *in vivo*. To get an impression of the specific impact of HM1 and HM2 and to exclude undesirable side effects that may arise from the luminal coiled-coil and SUN domains, we employed EGFP-tagged C-terminally truncated versions of the Sun4 full-length protein and analyzed their dynamic properties by FRAP. Not surprising, as shown in Fig. 5D, the C-terminal truncation led to a general overall increase in mobility compared to the full-length protein, which most likely can be ascribed to the disrupted coiled-coil and/or SUN domain interactions.

To effectively compare the different deletion constructs with each other, we have chosen two distinct reference values: 1) the averaged time-points when the respective constructs had regained 50% of their pre-bleach intensity (t_50%_-values) and 2) their relative intensity at 30 s post bleaching (Fig. 5E; a list with the detailed p-values can be found in Table S1). As can be seen in Fig. 5D-F, the construct containing both hydrophobic motifs, HM1 and HM2 (S4_ΔC-EGFP), featured the lowest mobility of the four constructs, with a t_50%_-value of 12.0 ± 2.4 s (mean ± s. d.) and 68.6 ± 4.9% (mean ± s. d.) signal recovery 30 s post-bleaching. The two constructs lacking either HM2 (S4_ΔHM2_ΔC-EGFP) or HM1 (S4_ΔHM1_ΔC-EGFP) both displayed significantly increased mobility compared to S4_ΔC-EGFP. S4_ΔHM1_ΔC-EGFP showed a t_50%_-value of 9.3 ± 2.1 s and reached 73.6 ± 5.5% of its pre-bleach intensity after 30 s. For S4_ΔHM2_ΔC-EGFP, the respective values were 7.1 ± 1.7 s and 73.8 ± 3.8. This reflects a rather similar degree of mobility as for S4_ΔHM2_ΔC-EGFP, since in this case only the t_50%_-value is significantly different each other (see Fig. 5E, F). The S4_ΔC_ΔHM1_ΔHM2-EGFP mutant lacking both hydrophobic motives showed by far the highest mobility, reaching 72.7 ± 9.0% of the initial signal intensity 1 s post-bleaching and nearly full recovery (97.1 ± 4.2%) within the first 30 s. This in other words means for HM1: (1) When HM1 is the only hydrophobic motif present in Sun4_ΔC, it reduces the protein’s mobility to a similar extent as HM2, the true transmembrane domain. (2) When the protein is factually integrated into the membrane (i.e. it contains HM2), the additional presence of HM1 provokes a weak, but -based on the parameters compared - discernible additional decrease in mobility (Fig. 5E, F). From this, we conclude that HM1, although not membrane-spanning, conveys NE targeting and - similar to the N-terminal hydrophobic motifs in Sun1 (Liu et al., 2007) - adds to the overall membrane retention of Sun4.

### Sun4 forms heteromeric assemblies with Sun3

Because full-length Sun4 appeared rather mobile when ectopically expressed in somatic NIH 3T3 cells, we hypothesized that in its natural context, i.e. in spermatids, Sun4 is hooked on spermatid-specific partners for its stabilization and retention within the NE.

Previous studies have shown that in round and elongating spermatids, Sun4 localizes to the regions where the MT manchette is in tight association with the NE (Calvi et al., 2015; Pasch et al., 2015). Notably, at these sites, Sun4 distributes rather congruently with Sun3. In addition, in previous *in vitro* experiments, we could co-immunoprecipitate Sun3 together with Sun4, indicating that these two proteins have an intrinsic capacity to interact with each other (Pasch et al., 2015). Therefore, it seems conceivable that in their natural environment, Sun4 and Sun3 are interacting partners forming joint LINC complexes, which in this combination are functional in a higher-order spermatid-specific nucleocytoplasmic network system.

To prove this hypothesis, we tested whether Sun4 indeed forms complexes with Sun3 in spermatids. In this regard, we performed co-immunoprecipitation (co-IP) from testicular cell suspensions of wild-type mice, using a protocol optimized for nuclear membrane proteins. In a first step, we employed 8 M urea extraction (see Lang et al., 1999), followed by high-speed centrifugation (100,000 x*g*) to selectively isolate the cellular membranes together with internal membrane proteins. Subsequently, the membranes were dissolved in RIPA buffer and subjected to classical co-immunoprecipitation using anti-Sun4 antibodies directed against the N-terminus of Sun4. In parallel, we used unspecific IgGs as control. Immunoprecipitates and corresponding supernatants were then probed with anti-Sun3 antibodies.

As expected, Sun4 was efficiently precipitated with the anti-Sun4 antibodies but was not detected in the control precipitate (Fig. 6A). Consistent with our hypothesis, Sun3 was also found in high amounts in the anti-Sun4 precipitate but not in the IgG control. This clearly demonstrates that Sun4 and Sun3 indeed interact with each other and form heteromeric assemblies not only in *in vitro* experiments but also in their natural background, i.e. in spermatids.

**Fig 6.**
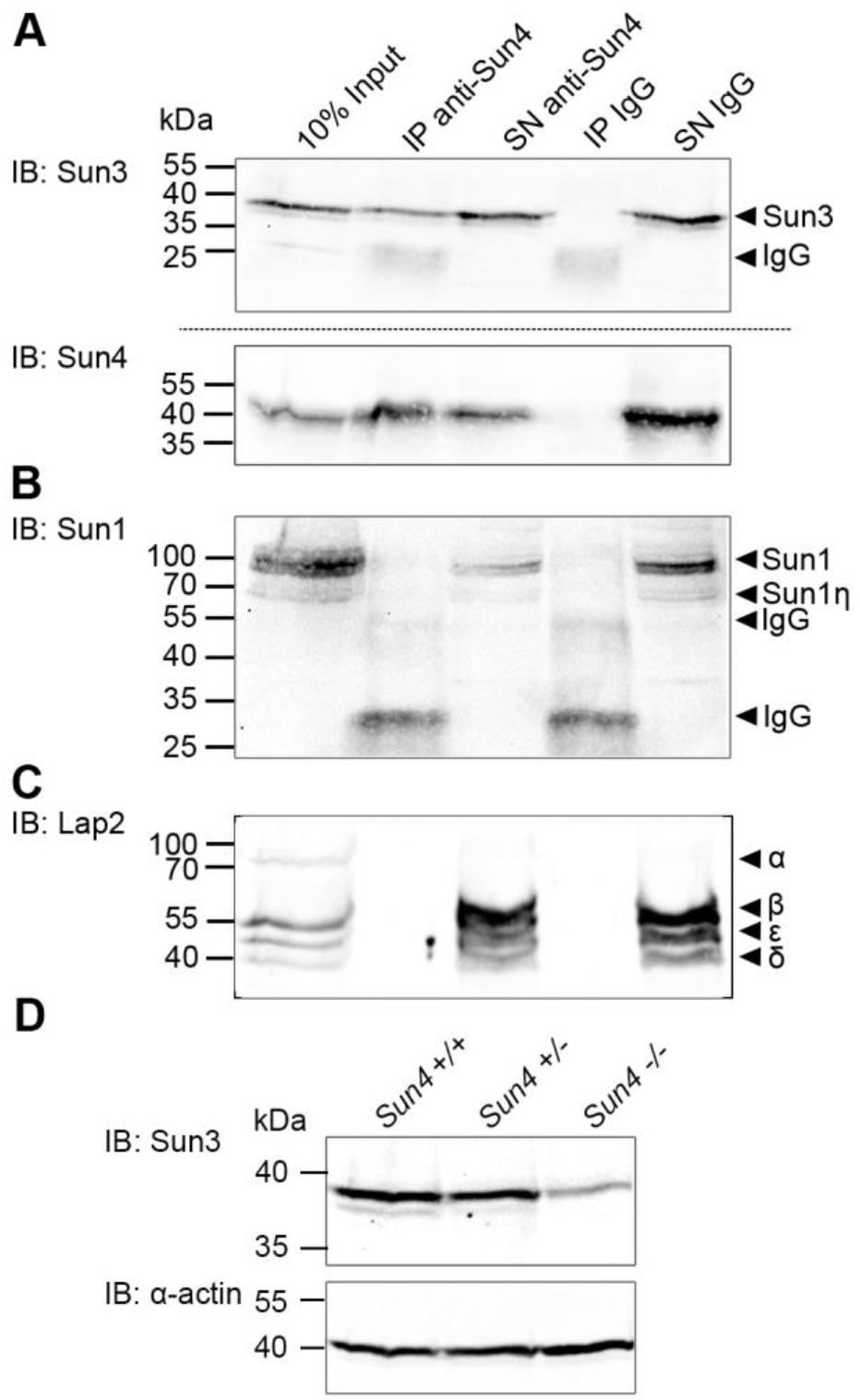
Sun4 forms heteromeric assemblies with Sun3 in mouse spermatids and Sun3 protein levels are decreased in Sun4 knock-out tissue. (A, B, C) Western Blot analyses of rabbit anti-Sun4 immunoprecipitated membrane protein extracts from mouse testis cell. Unspecific rabbit IgG was used as a negative control. Precipitates (IP) and supernatants containing unbound proteins (SN) were probed with (A) guinea pig anti-Sun3 and anti-Sun4, (B) anti-Sun1 and (C) mouse anti-Lap2 antibody. Untreated testis cells served as input control. Lap2 isoforms were assigned according to Berger et al., 1996. Please note that Lap2-α does not contain a TMD and thus can not be present in the SN. Amounts of loaded cell equivalents: 10% Input=1.5×10^6^; IP=1.5×10^7^; SN=1.5×10^7^. (D) Immunoblot detection of Sun3 levels in *Sun4^+/+^*, *Sun4^+/-^* and *Sun4^-/-^* testis tissue; α-actin protein levels served as loading control.

### Sun4 is required for maintenance of Sun3 expression levels

A recent study has shown that in spermatids, Sun3 protein considerably affects the expression of Sun4 in spermatids, i.e. Sun4 expression is downregulated in the absence of Sun3 (Gao et al., 2020). Therefore, we asked whether this is also true vice versa, i.e. whether Sun4 has a similar regulatory effect on Sun3 expression. To explore this, we studied the Sun3 protein levels in testis tissue samples of *Sun4^+/+^*, *Sun4^+/-^* and *Sun4^-/-^* mice using Western Blot analysis. As shown in Fig. 6D, in the *Sun4* knock-out condition, the amount of Sun3 appeared significantly reduced compared to the amounts in wild-type and heterozygous tissues, while the levels of α-actin expression, which was monitored as a control, remained comparable between all three genotypes. This finding suggests that Sun4 indeed is necessary to sustain Sun3 synthesis and that the two proteins thus might not only interact on a structural but also on a regulatory level *in vivo*.

### Sun4 interacts with Lamin B3 in mouse spermatids

In a previous study (Pasch et al., 2015), we demonstrated that depletion of Sun4 not only results in mislocalization of its direct LINC partners but also interferes with general NE integrity and leads to severe malformation of spermatids. Hence, it appears very likely that Sun4, directly or indirectly, is interconnected with other components of the NE, forming a nucleo-cytoplasmic interactome that, as a whole, orchestrates the process of sperm-head formation.

To explore whether Sun4 interacts with other constituents of the spermatid INM that show comparable posterior localization, i.e. Sun1 and Lap2, we probed Sun4 co-IP samples with antibodies against these two proteins. As shown in Fig. 6B, C, respectively, neither Sun1 nor Lap2 (or any of their isoforms) were detectable in the anti-Sun4 precipitate, indicating that none of the two proteins effectively binds to Sun4 or Sun4 containing complexes. On the other hand, the absence of Sun1 in the anti-Sun4 precipitate substantiates that the detected Sun4-Sun3 interaction described above is specific and not a result of a general intrinsic heteromerization tendency of SUN domain proteins, which could force interactions among one another *in vitro*.

Two other proteins that show a posterior localization during nuclear elongation are the peripheral NE proteins Lamin B1 and Lamin B3, the only two lamin isoforms expressed in spermatids (Schütz et al., 2005a; Vester et al., 1993). As described for Sun1 and Lap2, the distribution of the two Lamins as well partially overlaps with that of Sun3/Sun4.

Since SUN domain proteins as the INM LINC components are supposed to directly connect to the nuclear lamina (Starr and Fridolfsson, 2010), Lamins B1 and B3 are prime candidates for a direct interaction with Sun4.

To test whether Sun4 binds to one or both of them *in vivo*, we again performed Sun4 co-IP from testicular cells of wild-type mice. However, since Lamin B1 and Lamin B3 are nucleoplasmic proteins, we followed standard IP procedures and hence, directly subjected the cells to RIPA lysis. The resulting precipitates and supernatants were probed with both anti-Lamin B3 and anti-Lamin B1 antibodies.

Using this assay, Lamin B1 was not detectable in either the control lysate precipitated with unspecific IgG or the anti-Sun4 precipitate (Fig. S3), suggesting that it is not a direct interaction partner of Sun4. Lamin B3, however, as shown in Fig. 7B, indeed was detectable in the anti-Sun4 precipitate but not in the IgG control, a result that clearly supports the notion that Sun4 may directly or indirectly interact with Lamin B3 in spermatids.

**Fig 7.**
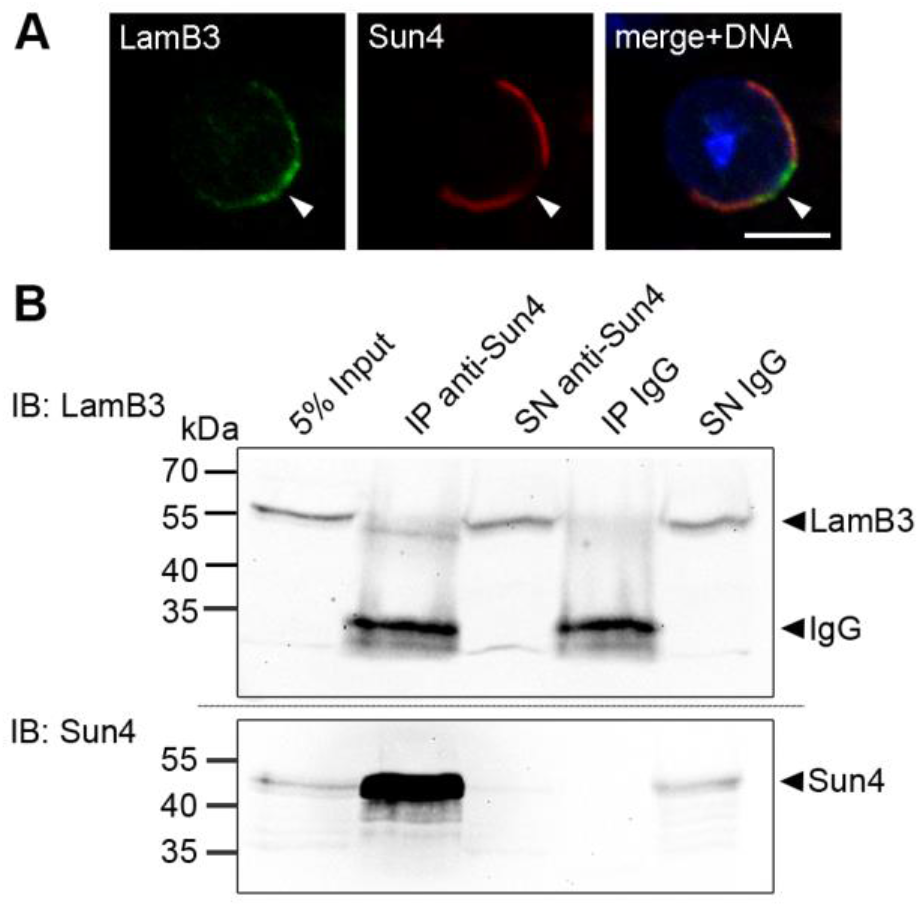
Sun4 binds to Lamin B3 in mouse spermatids. (A) Confocal microscopy image of an elongating spermatid co-immunolabelled with anti-Lamin B3 (LamB3; green) and anti-Sun4 antibodies (red). DNA was counterstained with Hoechst (blue). Scale bar: 5 µm. Arrowheads indicate the implantation fossa. (B) Immunoblot (IB) analysis of co-immunoprecipitation from mouse testicular suspension cells with guinea pig anti-Sun4 NTD antibody and unspecific guinea pig IgG as a negative control. Precipitates (IP) and supernatants containing unbound proteins (SN) were probed with rabbit anti-Lamin B3 and anti-Sun4 peptide antibodies. Untreated testis cells served as input control. Loaded cell equivalents: 5% Input=2×10^6^; IP=4×10^7^; SN=5×10^6^.

### Sun4 affects the distribution of Sun3 but not of the NE components Lamin B3, Lap2 and Lamin B1

In our previous study, we showed evidence that Sun4 depletion results in a severe mislocalization of its putative LINC complex partners Sun3 and Nesprin1. Moreover, in Sun4 deficient spermatids, we found a significantly altered distribution of Sun1/Nesprin3 compared to that found in wild-type cells (Pasch et al., 2015, see also Fig. 8A, A’).

**Fig 8.**
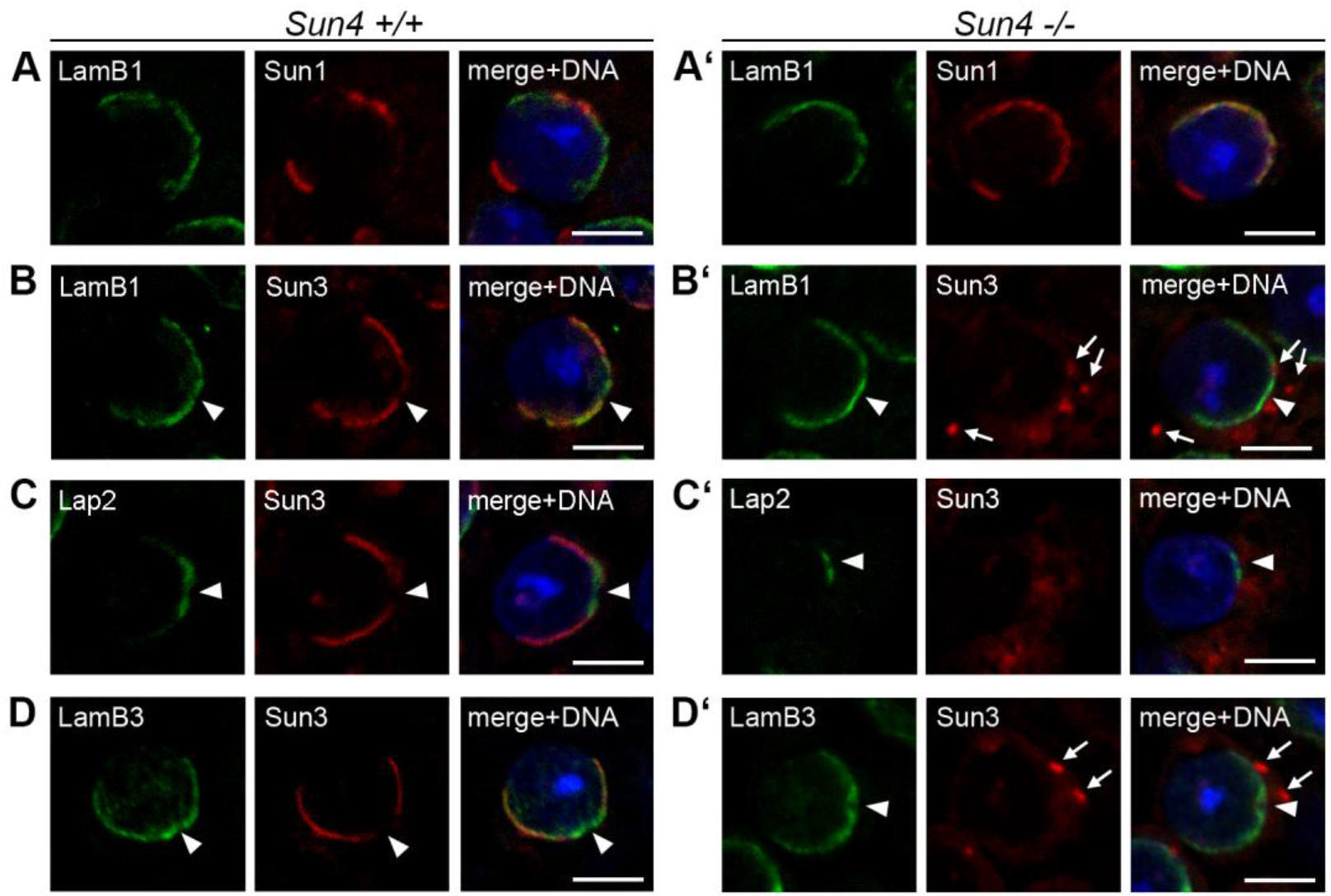
Sun4 deficiency affects Sun1, but not Lamin B3, Lamin B1 and Lap2 distribution. Testis paraffin sections of adult *Sun4^+/+^* and *Sun4^-/-^* mice were co-immunofluorescence-labelled with antibodies against (A, A’) Lamin B1 (LamB1; green) and Sun1 or (B, B’) Sun3 (red), (C, C’) Lap2 (green) and Sun3 (red) and (D, D’) LamB3 (green) and Sun3 (red). DNA was stained with Hoechst (blue). Shown are images of round and early elongating spermatids. Scale bars: 5 µm. Arrowheads point at the implantation fossa, arrows at cytoplasmic Sun3 aggregates typically found in *Sun4^-/-^* spermatids.

To explore to which extent depletion of Sun4 also affects the localization of other not yet analyzed NE components, we performed further co-immunofluorescence analysis on *Sun4^+/+^* and *Sun4^-/-^* testis tissue and tested for Sun3 distribution and, particularly, for Lap2, Lamin B1 and B3.

Consistent with earlier studies (Alsheimer et al., 1998; Göb et al., 2010; Vester et al., 1993), in a wild-type background, we found Lamin B1 and Lap2 in round spermatids situated at the posterior nuclear periphery, with a local concentration at the most posterior area neighboring the implantation fossa, and laterally overlapping with Sun3 (Figs. 8B, C). With progressing elongation, both proteins retreated more and more towards the posterior pole, a region where Sun3 is completely absent (as shown in Alsheimer et al., 1998 and Göb et al., 2010). On depletion of Sun4, Sun3 entirely disappeared from the nuclear periphery and accumulated outside the nucleus in spot-like aggregates as previously described (Fig. 8B’-D’; Pasch et al., 2015). Strikingly, absence of Sun4 did not result in any visible changes in the distribution patterns of Lamin B1 and Lap2. In the Sun4 deficient background, Lamin B1 and Lap2 still localized to the posterior NE. Contrary to the situation of Sun1, which in *Sun4^-/-^* spermatids caudally extends to the more lateral regions (Fig. 8A’; Pasch et al., 2015), their confinement to the posterior pole was apparently not affected either (Fig. 8B’, C’). In wild-type spermatids, Lamin B3, which we have identified as a *bona fide* Sun4 interaction partner (see above), also localized to the posterior NE has has been described previously (Schütz et al., 2005a). In addition, consistent with the previous findings, a significant proportion also localized to the nucleoplasm (Fig. 8D; Schütz et al., 2005a). Remarkably, in the *Sun4^-/-^* tissues, we were not able to detect any overt changes either the nucleoplasmic or the NE associated pool, which showed clear concentration around the fossa region (Fig. 8D’). Thus, although directly or indirectly interconnected with Sun4, the localization of Lamin B3 seems to be independent of Sun4.

## Discussion

Nuclear elongation and reorganization are key elements of male germ cell differentiation. Any defects in these processes usually lead to defective sperm function and thus cause severe fertility problems, which often results in entire male infertility (Yan, 2009). The exact mechanisms and networks underlying these typical shaping and restructuring events, however, are still poorly understood. According to current theories, sperm head formation and particularly nuclear elongation are guided by mechanical forces, which are supposed to be generated by highly organized cytoskeletal structures surrounding the nucleus, i.e. the acroplaxome and the manchette (reviewed Dunleavy et al., 2019 and Teves and Roldan, 2022).

In their role as nucleo-cytoskeletal linkers, LINC complexes represent perfect candidates for involvement in physically connecting these structures to the nucleus, thus enabling a transfer of cytoskeletal shaping forces (Dunleavy et al., 2019; Kmonickova et al., 2020; Kracklauer et al., 2010; Teves and Roldan, 2022). Consistently, in several previous studies, the spermiogenesis specific LINC complex components Sun4 and, very recently, Sun3 were identified as crucial players in sperm head formation, essential for proper NE attachment of the manchette microtubules, correct positioning of other LINC components and finally also for directed nuclear elongation (Calvi et al., 2015; Gao et al., 2020; Pasch et al., 2015; Yang et al., 2018). Although important for understanding its central function, the intrinsic molecular properties of Sun4 and its specific interplay with other components of the spermatid NE remained largely unknown. In particular, to date, it has remained unclear how Sun4 is oriented within the NE and how exactly it is organized within a functional LINC complex. In the present study, we have now tackled several of these remaining questions in detail.

### Sun4 shares the typical membrane topology of a SUN domain protein

Even though sharing the typical molecular composition of a SUN domain protein, the actual localization of Sun4 and its *de facto* orientation within the NE were not entirely clear. Several previous studies have stated scenarios for Sun4 that are in apparent conflict with the classical topology of SUN domain proteins (Shao et al., 1999; Tapia Contreras and Hoyer-Fender, 2021; Yang et al., 2012; Yang et al., 2018). A couple of years ago, based on a Sun4 immunogold-labeling experiment, Shao et al. (1999) showed that besides its nowadays accepted localization to the NE regions that are decorated by the manchette (Calvi et al., 2015; Gao et al., 2020; Pasch et al., 2015), Sun4 (= Spag4) appears to be also located within the axoneme. Using an *in vitro* assay, they found indications that within the axoneme, Sun4 may bind via a C-terminal leucine zipper motif to the cytoskeletal protein Odf1 (Shao et al., 1999). This, however, is difficult to reconcile with a classical SUN domain protein topology, as such an interaction would implicate a non-typical cytoplasmic orientation of the C-terminal SUN domain rather than a regular SUN domain typical localization within the perinuclear space (Crisp et al., 2006). Interestingly, bioinformatical tools predicted two hydrophobic, potentially transmembrane motifs within the Sun4 amino acid sequence, offering various possibilities of how Sun4 could be oriented within the NE - including those in which the SUN domain could actually locate to the cytoplasm. In such a scenario, when the SUN domain of Sun4 indeed is oriented to the cytoplasm, it may, hypothetically, then also directly bind to ODF1. Our present experiments, however, now definitely rule out this option. Our systematic biochemical screening, combined with immunogold labeling of the Sun4 NTD in spermatids, identified HM2 as the only factual TMD. Beyond this, we could provide unequivocal evidence that the C-terminal part of Sun4 locates in the PNS/ER compartment while the NTD is directed to the nuclear interior of the spermatid. Hence, we were now able to demonstrate that Sun4 actually shares a SUN domain protein typical membrane topology. Moreover, based on our present study, we now can rule out any direct interactions between the Sun4 CTD and any cytoplasmatic proteins. Also, we were unable to detect the previously described axonemal localization of SUN4 (Shao et al.,1999). Although we used antibodies from two different species and applied them on variably fixed testis-tissues, we could not reproduce a gold-labeling of the axoneme.

Instead, we could validate our previous observation that Sun4 is exclusively located within the NE, particularly enriched at the lateral posterior regions and the INM of the redundant nuclear envelope (Pasch et al., 2015; Figs 4, S1). It is worth noting that in recent studies, Yang et al. identified both, Odf1 and Sun4 as essential components for proper head-to-tail coupling (Yang et al., 2012; Yang et al., 2018). Considering these observations but also taking into account our results that Sun4 is *de facto* not present in the axoneme into account, we suggest that instead of operating as interacting binding partners at the axoneme, Sun4 and Odf1 could rather be part of a joint complex located at the rim of the basal plate, where it might have a yet largely unknown function in tethering the sperm tail to the nuclear envelope.

Our immunofluorescence and FRAP analyses suggest that the second hydrophobic element HM1, although not transmembrane, contributes to general membrane targeting and retention of Sun4. A similar phenomenon was previously described for Sun1. Liu et al. (2007) found that of the four HMs present within the molecular sequence of Sun1, only one is a true TMD, while the other three provide membrane affinity and thus add to the local retention of Sun1. How exactly this added value is mediated, however, remained largely unclear. Liu et al. (2007) suggested two possible scenarios that both, theoretically, might also apply for Sun4: (1) the additional hydrophobic elements could function as binding sites for other INM integrated proteins, or (2) they could interact directly with the INM lipid bilayer. In this context, a mechanism similar to that of nuclear lamins would be conceivable. Lamins are targeted and tethered to the INM by adding extra hydrophobic elements through post-translational modification, i.e. farnesylation of the CaaX box motif (Kitten and Nigg, 1991; for review, see Perez-Sala, 2007). However, differing from the lamins, in the case of the SUN proteins, the membrane targeting would be mediated by a short peptide motif instead of an isoprene group annex.

### Sun4 and Sun3 are mandatory LINC complex partners

In our FRAP analyses, we could also find that in a somatic context, compared to Sun1, Sun4 shows exceptional mobility within the NE. Hence, Sun4 overtly requires further, spermatid-specific binding partners for effective NE retention. Sun3 would be a good candidate for being one of these vital retention factors, as it has a congruent distribution, shares the capacity to bind Nesprin1 (Göb et al., 2010; Pasch et al., 2015), and, as shown in our current study, specifically co-precipitates with Sun4 from mouse spermatids. Interestingly, Gao and colleagues recently demonstrated that, vice versa, the Sun4 protein also co-precipitates with endogenous Sun3 (Gao et al., 2020). In their study, they also demonstrated that Sun4 localization to the NE implicitly depends on the presence of Sun3, a phenomenon that we could show several years before for Sun3 in a Sun4 deficient background. On Sun4 deficiency, Sun3 completely disappears from the spermatid NE and, vice versa, when Sun3 is absent, Sun4 is not recruited to the NE but locates to the cytoplasm, where it shows a diffuse distribution (Pasch et al., 2015; Calvi et al., 2015; Gao et al., 2020). Both, the Sun3 and Sun4 mouse models thus have provided clear proof for a strict interdependency of Sun3 and Sun4 regarding their localization and, hence, point to a coordinated function, either as parallel or as joint LINC complex assemblies. Remarkably, the NTD of Sun3 is pretty short and comprises only seven amino acids (Göb et al., 2010). Therefore, it appears rather unlikely that Sun3 is able to effectively anchor to nuclear structures Thus, Sun3 INM retention and correct NE localization somehow need support, e.g. by binding to Sun4, which in turn connects to the nuclear lamina. Therefore, we propose that Sun3 and Sun4 form joint LINC complexes or LINC complex systems, in which Sun3 stabilizes the connection to their putative KASH binding partner Nesprin1 and Sun4 provides the interconnection to the nuclear lamina.

The actual organization of such Sun3/Sun4/Nesprin1 LINC complexes, however, remains obscure. According to the current model, which is mainly based on structural analysis of the human SUN2-KASH1/2 complex, SUN proteins form coiled-coil trimers that connect to three KASH peptides (Sosa et al., 2012). It is however unclear, whether this also applies for Sun3 and Sun4, and if so, whether they form heteromeric coiled-coil timers or arrange into higher-order LINC systems as homotrimers. Structured illumination microscopy (Pasch et al., 2015) indicated that Sun3 and Sun4 may actually occur as homotrimers, but to some extent do also form joint heteromeric assemblies in the spermatid INM.

Interestingly, besides the overt above-described functional interdependency within the nucleo-cytoplasmic network system, the two proteins also seem to closely hinge on each other on a regulatory level. Recently, Gao et al. (2020) have shown that compared to the wild-type, the levels of Sun4 are significantly reduced in Sun3 depleted testis tissue. In our present study, we now show the reverse, i.e. that in a *Sun4^-/-^* background, the Sun3 protein expression is significantly reduced as well. So, apparently, both proteins require the presence of the other to keep their synthesis on a constant level. Considering the circumstance that, as described above, neither of the two proteins is functional on its own, a conceivable purpose of this reciprocal expression regulation could be that it might serve to keep the levels of the two proteins in balance and therefore prevent an accumulation of dysfunctional monomeric assemblies. This, however, currently is just speculation and to tackle this problem, further studies on this phenomenon need to be conducted in the future.

### Lamin B3 represents a *bona fide* nucleoskeletal binding partner of Sun4 containing LINC complexes

LINC complexes are classically defined as linkers of the nucleoskeleton and the cytoskeleton (Crisp et al., 2006). As evidenced by Sun3 and Sun4 knock-out models and corroborated by immunolocalization studies, Sun3/Sun4 containing LINC complexes connect to the MT manchette at the cytoplasmic surface of the NE (Calvi et al., 2015; Gao et al., 2020; Pasch et al., 2015). Respective nucleoskeletal counterparts, however, were not known so far. Somatic SUN domain proteins directly connect to the nuclear lamina with their NTDs (Starr and Fridolfsson, 2010). Since spermatids also form a nuclear lamina, though considerably modified regarding the basic composition and local distribution (Alsheimer et al., 1998; Schütz et al., 2005a; Vester et al., 1993), a similar scenario is conceivable for LINC complexes in spermatids as well. In spermatids, only two lamins, i.e. Lamin B1 and Lamin B3, are present (Schütz et al., 2005a; Vester et al., 1993). Thus, they represent the only possible lamin interaction partners for Sun3 and Sun4 Admittedly, in the case of Sun3, a direct interaction with the lamina appears pretty unlikely, due to its very short NTD (see above). The NTD of Sun4, however, consists of 165 amino acids (Fig. 1), which, according to our results, are directed to the nuclear interior (Fig. 4) and therefore are perfect target for direct interaction with the nuclear lamina.

Lamin B1 and B3 both locate to the posterior half of the nucleus in round spermatids, largely colocalizing with the Sun3/4 containing LINC complexes (Figs 7A, 8B). With progressing differentiation, the two lamins congregate at the very posterior pole, a region where Sun3 and Sun4 are actually not present. It is noteworthy that in the case of Lamin B3, a significant proportion is also distributed within the nucleoplasm, thus having no tight contact with the nuclear periphery and, hence, being no primary target for interaction with LINC complexes (Schütz et al., 2005a).

In an *in vitro* assay, Yeh et al. (2015) could find that human Lamin B1 co-precipitates with ectopically co-expressed Sept12 and Sun4 and that in this artificial context, these three proteins together can overtly form higher-order complexes. We, however, could not detect any hints for a direct or possibly indirect interaction between Sun4 and Lamin B1 in its natural context, i.e. spermatids. It has, admittedly, to be noted that Lamin B1 usually forms very stable, salt-resistant polymer networks (Schirmer and Gerace, 2004). Thus, with the RIPA buffer system, only a marginal portion of Lamin B1 could be solubilized in our *in vivo* co-IP assay and this modicum of soluble Lamin B1 may be too limited to detect true Sun4/Lamin B1 complexes. Therefore, we currently can not rule out that there is indeed no direct interaction between the Sun4 NTD and the endogenous Lamin B1 in spermatids.

However, in another experiment, we could identify Lamin B3 as a factual interaction partner of Sun4. Considering the differences in its local distribution compared to that of the Sun4 containing LINC complexes, we assume that of the entire Lamin B3 pool, only the fraction associated with the lateral NE regions can participate in such an interaction. In contrast to other B-type lamins, Lamin B3 appears to be highly dynamic, shuttling between the nucleoplasmic and peripheral pool. Lamin B3 is therefore assumed to support a flexibilization of the lamina network system in spermatids, which is required to facilitate the restructuring of the spermatid nucleus (Schütz et al., 2005b; Kracklauer et al., 2013). In the context of the striking, orchestrated nuclear envelope dynamics in spermatids that are crucial for nuclear shaping (for review, see Kracklauer et al., 2013), the specific interaction between Lamin B3 and Sun4 may hold a regulative function a) in transforming the cytoplasmic shaping forces to the nuclear interior and/or b) to support local nuclear envelope flexibilization by directing Lamin B3 to the posterior nuclear envelope, where it may weaken the basic rigid lamina scaffolding provided by Lamin B1. A comparable function was also postulated for meiosis-specific Lamin C2 in prophase I spermatocytes. Lamin C2 accumulates at the sites of telomere attachment, hence supporting meiotic telomere movements in an otherwise overall rigid lamina structure (Link et al., 2013).

Surprisingly, absence of Sun4 in spermatids did not result in severe effects on the general Lamin B3 distribution, indicating that the specific localization of Lamin B3 to the posterior NE is not exclusively dependent on Sun4 but may be backed decisively by other putative interaction partners such as the posterior localized Lap2 or Lamin B1 for example (Göb et al., 2010; Schütz et al., 2005a). Vice versa, Lamin B3 could also be critical for Sun4 localization, e.g. as a recruiting platform, and thus organize its function in connecting the MT manchette to the NE. Therefore, it would be very enlightening to investigate the distribution of Sun4 in a Lamin B3 deficient background in a future study.

### Sun4 functions at the nucleocytoplasmic junction

Previous studies have shown that Sun4 is essential for sperm head shaping and for correct assembly and NE attachment of the cytoplasmic MT manchette. Furthermore, upon Sun4 deficiency, its *bona fide* LINC complex partners, Sun3 and Nesprin1, entirely disappear from the NE, which demonstrates its crucial impact on the correct positioning of other NE components (Calvi et al., 2015; Pasch et al., 2015). Here, we continued investigating this specific role of Sun4 by searching for further Sun4 *in vivo* interaction partners and studying the effect of a Sun4 depletion on other constituents of the posterior NE. Besides the Lamins B1 and B3, which were extensively discussed above, Sun1 and Lap2 are two further NE components that are highly interesting in the context of nuclear shaping. Both, Sun1 and Lap2 (with the exception of Lap2α, which is a splice form that lacks a TM domain; see Vlcek et al., 1999) are integral proteins of the INM that, at least in the early steps of spermatid differentiation, are found located at the posterior NE where they occupy territories that are widely overlapping with that of Sun4 (Alsheimer et al., 1998; Göb et al., 2010). This offers the opportunity for direct or indirect interaction with Sun4, or rather renders the possibility of a mutual interrelationship that may impact the territorial distribution of the other NE components.

Our co-IP results demonstrated that neither Sun1 nor Lap2 can efficiently be precipitated together with Sun4 as bait, suggesting that they are not direct or even indirect binding partners of Sun4. Nonetheless, we previously identified Sun4 as an essential determinant for Sun1/Nesprin3 distribution (Pasch et al., 2015). In *Sun4^-/^*^-^ spermatids, we found the posterior Sun1/Nesprin3 signals “leaking” to the more lateral regions instead of being confined to the very posterior pole (Pasch et al., 2015; see also Fig. 8A, A’). Hence, Sun4 is apparently part of a kind of physical barrier for Sun1 containing LINC complexes, promoting their posterior confinement. However, this appears to apply only to Sun1 and its direct partners. In the case of Lamin B1 and B3 and also observed for Lap2, no overt change in localization was observed in a Sun4 deficient background, demonstrating that their distribution is rather independent of the presence of Sun4.

Notably, previous studies also disclosed that chromatin compaction in the spermatids is not vitally affected by Sun4 depletion (Calvi et al., 2015; Pasch et al., 2015). Since the posterior pole of the spermatid nucleus is the only site where the chromatin tightly contacts the NE (Collas and Poccia, 1996), and Lap2 and Lamin B1 are well-known to bind to chromatin, we suggest that the posterior confinement of Lap2, Lamin B1 and Lamin B3 may somehow be interrelated with the chromatin remodeling rather than being regulated by Sun4 containing LINC complexes. The INM protein Dpy19l2, which localizes to the anterior pole of the spermatid nucleus, might be an essential coordinator here: As shown by Pierre et al. (2012), in Dpy19l2 depleted spermatids, Lamin B1 loses its posterior polarization and extends to the more apical regions. Besides this, the acrosome and, remarkably, also the manchette fail to connect to the NE. This indicates that proper Sun4 localization and also the general LINC complex distribution and function may depend vitally on Dpy19l2 and its partners. Investigating these interrelations in more detail would be another interesting challenge for future studies.

In conclusion, our present study now provides clear evidence that Sun4 is an integral protein of the INM that shares a SUN domain protein typical type II membrane topology and, therefore, in all probability, can not directly interact with any of the cytoskeletal proteins. It exclusively locates to the lateral regions of the posterior spermatid NE, where it most likely forms heteromeric assemblies with Sun3. Via a specific interaction of the Sun4 NTD with Lamin B3, Sun4 anchors to the nuclear lamina and, together with Sun3, forms the INM constituents of spermatid-specific LINC complexes with Nesprin 1 as the *bona fide* KASH binding partner. Beyond this, we could also provide evidence that Sun4 is involved in regulating Sun3 expression. Thus, our results contribute to a better understanding of the Sun4 interactions and its function at the spermatid nucleocytoplasmic junction and the entire process of sperm-head formation.

## Material and Methods

### Ethics statement

All animal care and experimental protocols were performed according to the guidelines specified within the German Animal Welfare Act (German Ministry of Agriculture, Health and Economic Cooperation). Animal housing and breeding were approved by the local regulatory agency of the city of Würzburg (Reference ABD/OA/Tr; according to 111/1 No. 1 of the German Animal Welfare Act). All aspects of mouse work were carried out under strict guidelines to ensure careful, consistent, and ethical handling of the mice.

### Animals and tissue preparation

Testis tissue samples were obtained either from wild-type, heterozygous or knock-out litter-mates of the Sun4 knock-out mouse (*Mus musculus*) strain *(Spag4^tm1(KOMP)Mbp^*) (Pasch et al., 2015) or from wild-type mice of other C57BL/6J strains. Eight to sixteen weeks-old male mice were sacrificed using CO_2,_ followed by cervical dislocation. Testes were dissected and further processed for protein analysis or immuno-localization studies as described below.

### Generation of plasmid constructs

To obtain Myc-S4_FL-EGFP and Myc-S4_ΔC-EGFP, the complete Sun4 coding sequence (S4_FL) or the sequence coding for amino acids 1-195 (S4_ΔC) were amplified by PCR from the previously described full-length Sun4 cDNA (Pasch et al., 2015) using the sequence-specific primers Sun4_inc.ATG_5’ and Sun4_SUNdom_3’_woStop or Sun4_delC-term_3’ (Table S1) and inserted into cloning vector pSC-B (Agilent Technologies, Waldbronn, Germany). After sequence verification, the respective fragments were excised with EcoRI and ligated into pCMV-Myc (Clontech Laboratories, Mountain View, CA). To generate the constructs coding for the double tagged Sun4 peptide sequences, the resulting Myc-tagged versions were then cloned into the BamHI site of pEGFP-N3 (Clontech Laboratories). Finally, to eliminate an undesired TAG-codon that was introduced with the cloning procedure between the 5’ Myc-tag sequence and the Sun4 coding region, we performed site-directed mutagenesis-PCR to exchange the TAG by CAG (primers: Muta_Myc-S4_5’ and Muta_Myc-S4_3’; see Table S1).

Myc-S4_ΔHM1_ΔC-EGFP, Myc-S4_ΔHM2_ΔC-EGFP, Myc-S4_ΔHM1_ΔHM2_ΔC-EGFP were generated from the S4_ΔC-pCMV-Myc construct by deletion PCR using primers Sun4_delTM1_3’, Sun4_TM2_5’ (for deletion of HM1) or Sun4_CC_5’, Sun4_delTM2_3’ (for deletion of HM2) (see Table S2).

For generating S4_FL-EGFP and S4_ ΔC-EGFP, the respective Sun4 fragments were cut out from Myc-S4-pEGFP-N3 or Myc-S4_ΔC-pEGFP-N3 and cloned into the EcoRI-site of pEGFP-N2. EGFP-tagged HM deletion constructs were generated from S4_ ΔC-EGFP as described above.

To generate S1_FL-EGFP, the Sun1 coding sequence was amplified from Sun1 full-length cDNA (Göb et al., 2010) with primers Sun1_5’ and Sun1_3’_woStop (Table S2) and cloned into the SmaI site of pEGFP-N2.

### Cell culture and transfection

For our experiments, we used COS-7 (DSMZ; ACC-60) and NIH/3T3 (ATCC; CRL-1658). Cells were grown in DMEM (Invitrogen) supplemented with 10% FCS and 1% penicillin/streptomycin at 37°C and 5% CO_2_.

COS-7 cells were transfected with the respective Myc-Sun4-pEGFP fusion constructs using Effectene^TM^ (Qiagen) following the manufacturer’s protocol and incubated overnight before analysis of the ectopically expressed fusion proteins.

Transfection of 3T3 cells was carried out using the Matra^TM^ magnetic bead transfection system (IBA Lifesciences) to warrant nuclear membrane integrity for subsequent FRAP experiments. To enhance transfection efficiency, 3T3 cells were treated twice in a row with the respective construct in the prepared transfection mixture.

Before analysis, cells were further incubated for approx. 48 hours at 37°C and 5% CO_2_.

### Antibodies

Primary antibodies used in this study were: guinea pig and rabbit anti-Sun4, guinea pig anti-Sun3, guinea pig anti-Sun1, rabbit anti-LaminB1, and rabbit anti-LaminB3, mouse anti-Lap2, rabbit anti-LaminA/C, mouse anti-α-actin, mouse anti-PDI RL90, mouse anti-GFP (B-2) and mouse anti-Myc.

Secondary antibodies were: AlexaFluor 488 goat anti-guinea pig, AlexaFluor 488 goat anti-rabbit, Texas red goat anti-mouse, horseradish peroxidase goat anti-guinea pig, - rabbit or -mouse, and 6 nm gold donkey anti-guinea pig and goat anti-rabbit.

Detailed information about sources and working dilutions of the antibodies used in this study is listed in Tables S3 and S4.

### Immunocytochemistry

#### Immunofluorescence labeling of tissue sections

Halved testes were fixed in 1% PBS buffered FA for three hours at room temperature and further processed for paraffin embedding as described in Alsheimer et al., 2005. Embedded tissues were cut into 3-6 µm sections using a paraffin microtome (Leitz) and placed on SuperFrost^®^ Plus slides (Menzel Gläser, Thermo Scientific). Deparaffination and antigen retrieval was done as described previously (Link et al., 2016). Specimen were permeabilized with 0.1% Triton X-100 in PBS for 10 minutes, washed in PBS and blocked with PBT (0.15% BSA, 0.1% Tween 20 in PBS) for 1 hour. The samples were incubated with the appropriate primary antibodies (dissolved in PBT) for one hour at room temperature or, depending on the antibody, overnight at 4°C, washed with PBS and subjected to corresponding fluorescently labeled secondary antibodies (dissolved in PBS) for 30 minutes. DNA was counterstained with Hoechst 33258 (Serva).

#### Immunofluorescence labeling of transfected culture cells

Transfected cells were fixed with 1% FA in PBS for 3 minutes, permeabilized with 0.1% Triton X-100 in PBS for 5 minutes and blocked for 90 minutes with PBT at room temperature. Incubation with primary and secondary antibodies was done as described above.

#### Immunogold labeling

Immunogold labeling was done either on sections of shock frozen native testis-tissue or on sections of pre-fixed TissueTek^®^ O.C.T Compound embedded (Sakura) testes. For embedding in TissueTek^®^ O.C.T Compound, dissected testes were cut into halves and fixed for one hour at room temperature in 1% PBS buffered FA, followed by two washing steps in PBS for 30 minutes each. The fixed tissue was then dehydrated by consecutive soaking in 15% and 30% sucrose solution (in PBS), embedded in TissueTek^®^ O.C.T Compound and frozen in precooled (−140°C) methyl butane. The frozen tissue was sectioned to 7-10 µm using a 2800 Frigocut E cryo-microtome (Reichert-Jung; Leica Instruments, Nussloch, Germany). Sections of the native tissue were fixed in 1% FA (in PBS). Fixed tissue of both types was permeabilized with 0.05% Triton X-100 (in PBS) for 8-10 minutes, washed in PBS and blocked with PBT for 60-90 minutes at room temperature. Incubation with primary antibodies was done with anti-Sun4 rabbit antibody overnight at 4°C or with Sun4 guinea pig antibody for 1 hour at room temperature. Following washing in PBS, samples were subjected to corresponding gold-conjugated secondary antibodies for 2 hours at room temperature (anti-rabbit) or overnight at 4°C (anti-guinea pig).

### *In situ* proteinase K digestions

*In situ* proteinase K digestions were done according to Liu et al., 2007. Cos7 cells were transfected with Myc-/EGFP-tagged Sun4 constructs (four 35-mm dishes for each construct). 24 hours after transfection, the cells were washed twice with ice-cold KHM buffer (110 mM KOAc, 2 mM MgCl_2_, 20 mM HEPES, pH 7.4) for one minute each. One well per construct was incubated with 4 µg/ml proteinase K in KHM, one with 4 µg/ml proteinase K and 0.5% TritonX-100 in KHM and a third one in KHM without any additives, each for 45 minutes at room temperature. A fourth well of each construct was first incubated with 24 µM ice-cold digitonin in KHM for 10 minutes, washed in KHM, and subsequently digested in 4 µg/ml proteinase K solution for 45 minutes. After digestion, 40 µg/ml PMSF and 1:100 protease inhibitor mix (stock solution: 5 mM α-aminocaproic acid, 1 mM benzamidine, 1 mM EDTA, 2 μg/ml aprotinin, 2 μg/ml antipain, 2 μg/ml chymostatin, 2 μg/ml leupeptin, 2 μg/ml pepstatin) were added. Cells were transferred into reaction tubes, washed in KHM, and boiled in 2x SDS-sample buffer (120 mM Tris/HCl, 10% SDS, 20% glycerol, 20% 2-mercaptoethanol, bromophenol blue, pH 6.8) at 95°C for 5-10 minutes. The samples were separated by SDS-PAGE, transferred to nitrocellulose membranes and probed with anti-Myc and anti-GFP antibodies.

### Co-Immunoprecipitation

Co-IP experiments were performed with testicular suspension cells of nine to sixteen weeks-old mice. For co-IP analysis of LaminB3 with Sun4, we lysed the cells in RIPA buffer (1% Triton X-100, 0.5% sodium deoxycholate, 0.1% SDS, 1 mM β-glycerophosphate, 1 mM Na_3_VO_4_, 1 mM EDTA, 1 mM EGTA, in PBS) supplemented with 1x proteinase inhibitor cocktail (#04693132001; Roche) for 20 minutes on ice with occasional inverting. The lysate was cleared by centrifugation at 14,000 x*g* and 4°C. The supernatant was divided into two equal aliquots, and each aliquot was incubated with either guinea-pig anti-Sun4 antibody or unspecific guinea pig IgG (1 µg antibody per approx. 1×10^8^ cell equivalents each), roll-shaking over-night. Immuno-complexes were pulled with Dynabeads^TM^ Protein G (10003D, Invitrogen^TM^ by Thermo Fisher Scientific): 1.0 mg RIPA-pre-equilibrated beads per approx. 1×10^8^ cell equivalents were added to each aliquot, and the lysate-bead mixtures were roll-shaken for 2 hours at 4°C.

The protein-loaded beads were harvested using a magnet and washed three times with 250 µl RIPA buffer. The supernatant was used as a control for pull-down efficiency. Proteins contained in the supernatant were precipitated with acetone according to standard procedures. Precipitated proteins and immunocomplex decorated beads were resuspended in 2x SDS-sample buffer and heated to 95°C for 15 minutes. Denatured samples were analyzed by SDS-PAGE and Western blotting. For detection of interactions of Sun4 with other transmembrane proteins, we implemented the following adaption: Prior to co-IP, cellular membranes and membrane-associated proteins were extracted from approx. 2×10^8^ cells by lysis in 5 ml urea buffer (8 M urea, 100 mM Tris, 1 mM DTT; pH 8.0) for 20 minutes at room temperature. The lysate was centrifuged at 100,000 *x*g and 4°C for 1 hour to precipitate the membranes. The resulting pellet, including the integral membrane proteins, was repeatedly washed with PBS supplemented with protease inhibitor cocktail and reconcentrated using an Amicon^®^ Ultra-0.5 device with 10 K MWCO (Merck) until residual urea was diluted below 10 mM. Then the pellet was steeped in 2 ml RIPA buffer at 4°C overnight with continuous roll-shaking to extract protein complexes from the membranes. Insoluble pellet fragments were removed by centrifugation at 15,000 x*g* and 4°C. Dissolved immuno-complexes were collected as described above, using 2 µg of rabbit anti-Sun4 antibody and unspecific rabbit IgG, respectively, per approx. 1×10^8^ cell equivalents.

### SDS-PAGE and Western blotting

Protein samples were resuspended in 2x SDS-sample buffer, denatured at 95°C for several minutes and separated by SDS-PAGE according to standard procedures. For Western Blotting, proteins were transferred to nitrocellulose membranes. Membranes were blocked in TBST buffer (10 mM Tris/HCl, 150 mM NaCl, 0.1% Tween 20, pH 7.4) supplemented with 10% dry milk powder overnight at 4°C and incubated for 60-90 minutes at room temperature with respective primary antibodies in blocking solution. After three 10 minutes washes in TBST buffer, the membranes were incubated with corresponding peroxidase-coupled secondary antibodies. Bound antibodies were detected with chemiluminescent substrate (Western Lightning^®^ Plus, Perkin Elmer) and visualized with iBright^TM^ CL1000 (Thermo Fisher Scientific).

### Fluorescence microscopy and FRAP

Fluorescence images were acquired on a Leica TCS-SP2 AOBS confocal laser scanning microscope with a 63×/1.40 HCX PL APO oil-immersion objective and a pinhole setting of 1.0 P AU (Leica Microsystems, Wetzlar, Germany). Images shown are maximum projections of three sequential images. Images were processed with ImageJ (http://imagej.nih.gov/ij) and Adobe Photoshop CS5 (Adobe Systems, San Jose, CA).

For FRAP experiments, the transiently transfected cells were analyzed in FluoroDish culture dishes (World Precision Instruments, Sarasota, FL) using the 488 nm laser line of an Ar/Kr laser with pinhole setting 1.6 P AU. Fluorescence emission of the EGFP protein was detected at 494–563 nm. FRAP measurements were performed according to Jahn et al., 2010. Two single pre-bleach scans were acquired, followed by eight bleach pulses (100% laser intensity) within a circular bleach spot of 2 µm diameter. Recovery data were collected from single section images with 2x accumulation at 1 second (30 images), 2 seconds (30 images) and 5 seconds intervals (35 images). For imaging, the laser power was set to 7% of bleach intensity. Resulting intensity values of fluorescence emission were background-subtracted according to Phair and Misteli, 2000, using LCS Leica Confocal Software (Copyright 1997-2004 by Leica Microsystems). Corresponding FRAP curves were generated in GraphPad Prism version 9.0.1 (and following) for Windows (GraphPad Software, San Diego, California, USA). Presented recovery values are means ± s.d. of 10-15 different cells originating from at least five different experiments. Time points at 50% signal recovery (t_50_ values) were calculated as follows:

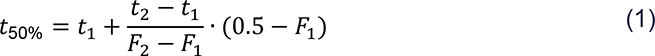

With *t*_50%_ the time of 50% fluorescence recovery, *t*_1_ the last timepoint before 50% recovery, *t*_2_ the first timepoint after 50% recovery, *F*_1_ the fluorescence intensity at *t*_1_, *F*_2_ the fluorescence intensity at *t*_2_.

Statistical analysis, including tests for statistical significance, was done with one-way ANOVA followed by Tukey’s multiple comparisons test using GraphPad Prism version 9.0.1 (and following). Corresponding whisker plots were also generated with GraphPad Prism.

### Electron microscopy

Immunogold-labelled testes sections were fixed in 2.5% glutaraldehyde (2.5% glutaraldehyde, 50 mM KCl, 2.5 mM MgCl, 50 mM cacodylate; pH 7.2) for 45 minutes, subsequently washed in 50 mM cacodylate buffer (pH 7.2) and post-fixed with 1-2% osmium tetroxide in 50 mM cacodylate for 1 hour on ice. Following several washing steps in H_2_O, the samples were dehydrated on ice in an increasing ethanol series, incubated two times in propylene oxide for 5 minutes each, and embedded in Epon as described in Link et al., 2016. Ultrathin tissue sections (50-60 nm) were cut on a Leica EM UC7 ultramicrotome and transferred onto copper grids (50 mesh). Sections were stained with uranyl acetate and lead citrate according to standard procedures. Samples were analyzed with a JEOL JEM-2100 transmission electron microscope operated at 200 kV (Jeol, Eching, Germany).

## Acknowledgments

We thank Silke Braune, Cornelia Heindl, Isabell Köblitz and Elisabeth Meyer-Natus for excellent technical assistance and Monika Leubner for support in generating S1_FL-EGFP. We are grateful to Christian Stigloher and his team (Imaging Core Facility, Biocenter, University of Würzburg, Germany) for helpful advice in EM sample preparation and electron microscopy, to Ulrike Kutay and Michael Reil (ETH, Zürich, Switzerland) for fruitful discussions and we are grateful to Nicola Jones for proofreading the manuscript.

---

**Figure S1. The Sun4 N-terminal domain is localized at the RNE of elongating spermatids.**

Representative electron micrographs of elongating spermatids in native mouse testis cryo-sections with immunogold-labelling (6 nm colloidal gold) of the Sun4 NTD. Area A is enlarged in A’. Gold particles (arrows) are mainly localized along the inner membrane of the redundant nuclear envelope and to both sides of the fossa region, but not directly within the implantation fossa (A’). Scale bars: 1 µm (A) and 200 nm (A’). Bb, basal body; Nu, nucleoplasm; Ac, acrosome; F, implantation fossa; RNE, redundant nuclear envelope, dc, distal centriole; pc, proximal centriole.

**Figure S2. Subcellular localization of EGFP-tagged Sun1 and Sun4 reporter constructs used for FRAP analysis.** EGFP-tagged full-length Sun1 and Sun4, as well as Sun4 constructs with deletion of the C-terminal coiled-coil and SUN domain were transiently transfected into NIH 3T3 cells. Confocal microscopy was used to monitor their expression patterns 48 h after transfection via their EGFP-tag. For each construct, a fluorescence image of one representative cell is shown. DNA was counterstained with Hoechst. Scale bars: 10 µm.

**Figure S3. Sun4 does not appear to bind to Lamin B1 *in vivo.*** Mouse testicular suspension cells were lysed in RIPA buffer and subjected to co-immunoprecipitation with guinea pig anti-Sun4 NTD antibody and unspecific guinea pig IgG as a negative control. The precipitates (IP) and supernatants containing unbound proteins (SN) were analyzed by immunoblotting (IB) using a rabbit anti-Lamin B1 antibody. The corresponding Sun4 control can be found in Fig. 6, as the same co-IP samples were used for both the Lamin B1 and the Lamin B3 assays. Untreated testis cells served as input control. Amounts of loaded cell equivalents: 5% Input=2×10^6^; IP=4×10^7^; SN=5×10^6^.

**Table S1: Details on statistical analysis of FRAP experiments (Output of GraphPad Prism multiple comparisons analysis)**

**Table S2: Oligonucleotides used in this study.**

**Table S3: Primary antibodies used in this study. IF, immunofluorescence; EM, electron microscopy (immunogold labeling); IB, immunoblot; mAb, monoclonal antibody; pAb, polyclonal antibody.**

**Table S4: Secondary antibodies used in this study.**

