## Supplemental Material for "Sun4 is a type II transmembrane protein of the spermatid inner nuclear membrane that forms heteromeric assemblies with Sun3 and interacts with Lamin B3"

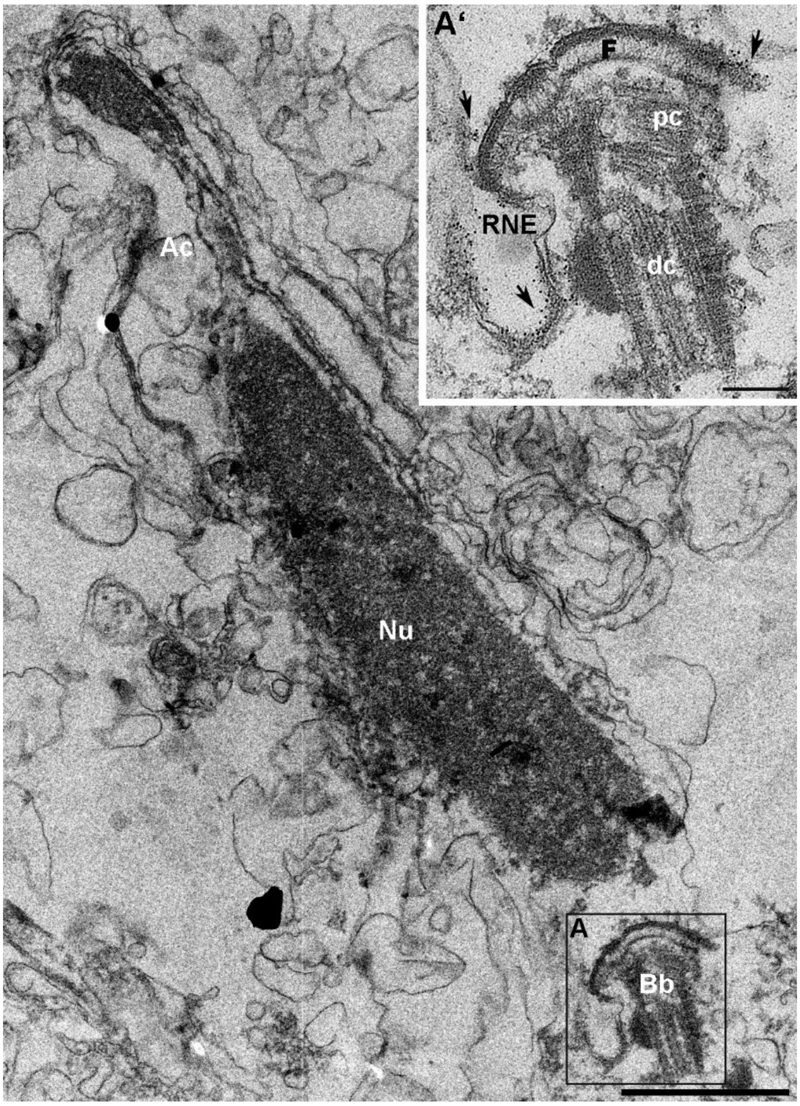


**Figure S1. The Sun4 N-terminal domain is localized at the RNE of elongating spermatids.**

Representative electron micrographs of elongating spermatids in native mouse testis cryo-sections with immunogold-labelling (6 nm colloidal gold) of the Sun4 NTD. Area **A** is enlarged in **A'**. Gold particles (arrows) are mainly localized along the inner membrane of the redundant nuclear envelope and to both sides of the fossa region, but not directly within the implantation fossa (A'). Scale bars: 1 µm (A) and 200 nm (A'). Bb, basal body; Nu, nucleoplasm; Ac, acrosome; F, implantation fossa; RNE, redundant nuclear envelope, dc, distal centriole; pc, proximal centriole.

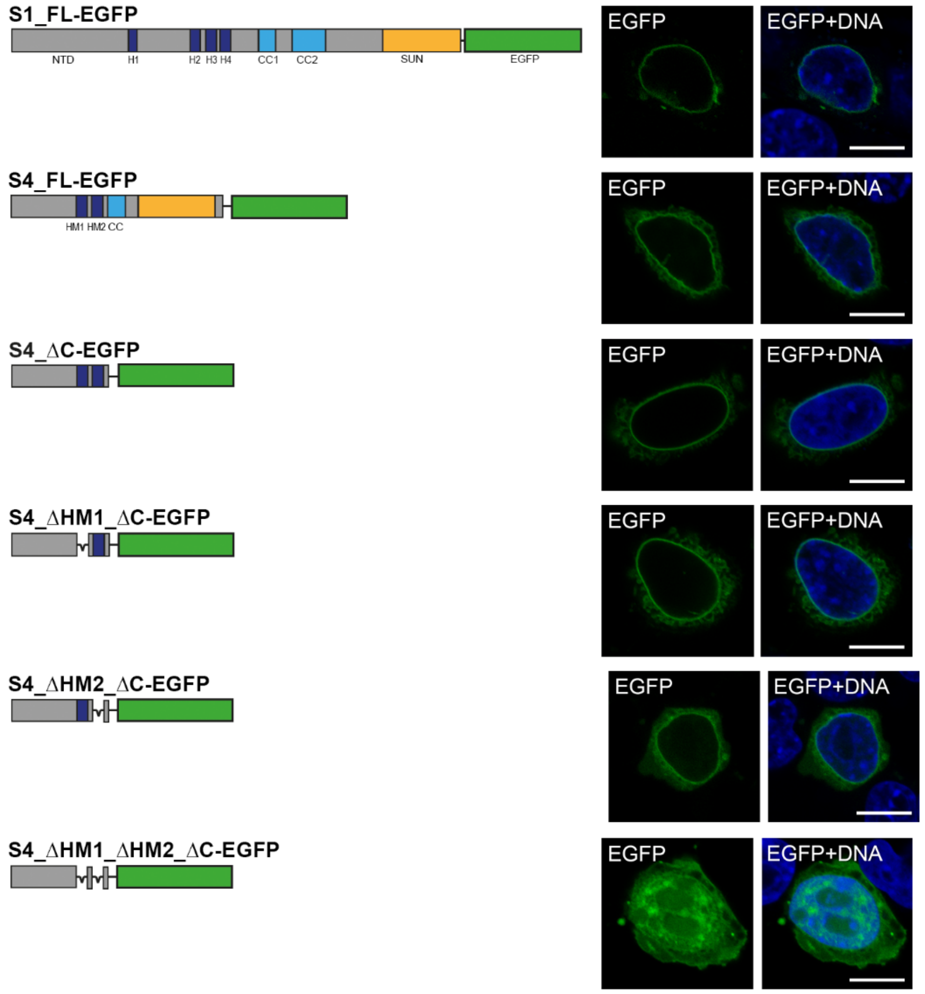


**Figure S2. Subcellular localization of EGFP-tagged Sun1 and Sun4 reporter constructs used for FRAP analysis.** EGFP-tagged full-length Sun1 and Sun4, as well as Sun4 constructs with deletion of the C-terminal coiled-coil and SUN domain were transiently transfected into NIH 3T3 cells. Confocal microscopy was used to monitor their expression patterns 48 h after transfection via their EGFP-tag. For each construct, a fluorescence image of one representative cell is shown. DNA was counterstained with Hoechst. Scale bars: 10 µm.

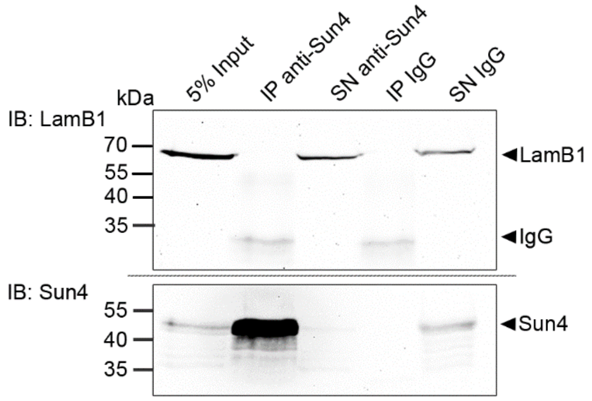


**Figure S3. Sun4 does not appear to bind to Lamin B1 *in vivo.*** Mouse testicular suspension cells were lysed in RIPA buffer and subjected to co-immunoprecipitation with guinea pig anti-Sun4 NTD antibody and unspecific guinea pig IgG as a negative control. The precipitates (IP) and supernatants containing unbound proteins (SN) were analyzed by immunoblotting (IB) using a rabbit anti-Lamin B1 antibody. The corresponding Sun4 control can be found in Fig. 6, as the same co-IP samples were used for both the Lamin B1 and the Lamin B3 assays. Untreated testis cells served as input control. Amounts of loaded cell equivalents: 5% Input=$2x10^6$; IP=$4x10^7$; SN=$5x10^6$.

### Table S1: Details on statistical analysis of FRAP experiments (Output of GraphPad Prism multiple comparisons analysis)

#### Ordinary one-way Anova of t50%-values - Multiple Comparisons

| | |
|---|---|
| Number of families | 1 |
| Number of comparisons per family | 3 |
| Alpha | 0.05 |

| **Tukey's multiple comparisons test** | Mean Diff. | 95.00% CI of diff. | Below threshold? | Summary | Adjusted P Value |
|---|---|---|---|---|---|
| S4_ΔC-EGFP vs. S4_ΔHM1_ ΔC-EGFP | 2.745 | 0.8906 to 4.99 | Yes | ** | 0.0024 |
| S4_ΔC-EGFP vs. Sun4_ΔHM2_ ΔC-EGFP | 4.903 | 3.049 to 6.757 | Yes | **** | <0,0001 |
| S4_ΔHM1_ ΔC-EGFP vs. Sun4_ΔHM2_ ΔC-EGFP | 2.158 | 0.3042 to 4.013 | Yes | * | 0.0192 |

| **Test details** | Mean 1 | Mean 2 | Mean Diff. | SE of diff. | n1 | n2 | q | DF |
|---|---|---|---|---|---|---|---|---|
| S4_ΔC-EGFP vs. S4_ΔC_ ΔHM1-EGFP | 12.03 | 9.289 | 2.745 | 0.7632 | 15 | 15 | 5.086 | 42 |
| S4_ΔC-EGFP vs. Sun4_ΔC_ ΔHM2-EGFP | 12.03 | 7.131 | 4.903 | 0.7632 | 15 | 15 | 9.086 | 42 |
| S4_ΔC_ ΔHM1-EGFP vs. Sun4_ΔC_ ΔHM2-EGFP | 9.289 | 7.131 | 2.158 | 0.7632 | 15 | 15 | 4 | 42 |

#### Ordinary one-way Anova of fluorescence intensities 30 s post-bleaching - Multiple Comparisons

| | |
|---|---|
| Number of families | 1 |
| Number of comparisons per family | 6 |
| Alpha | 0.05 |

| **Tukey's multiple comparisons test** | Mean Diff. | 95.00% CI of diff. | Below threshold? | Summary | Adjusted P Value |
|---|---|---|---|---|---|
| S4_ΔC-EGFP vs. S4_ΔHM1_ΔC-EGFP | -0.0502 | -0.09558 to -0.004812 | Yes | * | 0.0247 |
| S4_ΔC-EGFP vs. S4_ΔHM2_ΔC-EGFP | -0.05253 | -0.09791 to -0.007142 | Yes | * | 0.0172 |
| S4_ΔC-EGFP vs. S4_ΔHM1_ΔHM2_ΔC-EGFP | -0.2849 | -0.3330 to -0.2367 | Yes | **** | <0,0001 |
| S4_ΔHM1_ΔC-EGFP vs. S4_ΔHM2_ΔC-EGFP | -0.00233 | -0.04771 to 0.04305 | No | ns | 0.9991 |
| S4_ΔHM1_ΔC-EGFP vs. S4_ΔHM1_ΔHM2_ΔC-EGFP | -0.2347 | -0.2828 to -0.1865 | Yes | **** | <0,0001 |
| S4_ΔHM2_ΔC-EGFP vs. S4_ΔHM1_ΔHM2_ΔC-EGFP | -0.2323 | -0.2805 to -0.1842 | Yes | **** | <0,0001 |

| **Test details** | Mean 1 | Mean 2 | Mean Diff. | SE of diff. | n1 | n2 | q | DF |
|---|---|---|---|---|---|---|---|---|
| S4_ΔC-EGFP vs. S4_ΔHM1_ΔC-EGFP | 0.686 | 0.7362 | -0.0502 | 0.01711 | 15 | 15 | 4.149 | 53 |
| S4_ΔC-EGFP vs. S4_ΔHM2_ΔC-EGFP | 0.686 | 0.7385 | -0.05253 | 0.01711 | 15 | 15 | 4.341 | 53 |
| S4_ΔC-EGFP vs. S4_ΔHM1_ΔHM2_ΔC-EGFP | 0.686 | 0.9708 | -0.2849 | 0.01815 | 15 | 12 | 22.2 | 53 |
| S4_ΔHM1_ΔC-EGFP vs. S4_ΔHM2_ΔC-EGFP | 0.7362 | 0.7385 | -0.00233 | 0.01711 | 15 | 15 | 0.1926 | 53 |
| S4_ΔHM1_ΔC-EGFP vs. S4_ΔHM1_ΔHM2_ΔC-EGFP | 0.7362 | 0.9708 | -0.2347 | 0.01815 | 15 | 12 | 18.29 | 53 |
| S4_ΔHM2_ΔC-EGFP vs. S4_ΔHM1_ΔHM2_ΔC-EGFP | 0.7385 | 0.9708 | -0.2323 | 0.01815 | 15 | 12 | 18.1 | 53 |

**Table S2: Oligonucleotides used in this study.**

| Primer | Sequence | Purpose |
|---|---|---|
| **Sun4_inc.ATG_5’** | 5‘-AGGTCAGGATGCGGCGGA-3‘ | Forward primer for amplification of Sun4 and SUN4_ΔCTD cDNAs to create entry StrataClone construct |
| **Sun4_SUNdom_3’_woStop** | 5‘-ATGGGGTCCCCCTGTGA-3‘ | Reverse primer for amplification of Sun4_FL cDNA to create StrataClone construct |
| **Sun4_delC-term_3’** | 5‘-CAGCATCTCCGTAGGTTCGTT-3‘ | Reverse primer for amplification of SUN4_ΔCTD cDNA to create StrataClone construct |
| **Sun4_delTM1_3’** | 5‘-CCTCGGGGTAGGCATCTC-3‘ | Flanking primers for deletion of HM1 |
| **Sun4_TM2_5'** | 5‘-TGCAGGGAAATCTGCTCC-3‘ | |
| **Sun4_delTM2_3'** | 5‘-GGAGCAGATTTCCCTGCA-3‘ | Flanking primers for deletion of HM2 |
| **Sun4_CC_5'** | 5‘-CCTTTGGAGAACGAACCTACG -3‘ | |
| **Muta_Myc-S4_5’** | 5’-CCGAATTCGCCCTCAGGTCAGGATGC-3’ | Forward primer for site-directed mutagenesis to eliminate undesired TAG-codon |
| **Muta_Myc-S4_3’** | 5’- GCATCCTGACCTGAGGGCGAATTCGG-3‘ | Reverse primer for site-directed mutagenesis to eliminate TAG-codon |
| **Sun1_5’** | 5’-ATGGACTTTTCTCGGCTGCAC-3’ | Forward primer for amplification of Sun1 coding sequence |
| **Sun1_3’_woStop** | 5’-CTGGATGGGCTCTCCGTG-3’ | Reverse primer for amplification of Sun1 coding sequence |

**Table S3: Primary antibodies used in this study.** IF, immunofluorescence; EM, electron microscopy (immunogold labeling); IB, immunoblot; mAb, monoclonal antibody; pAb, polyclonal antibody.

| Antibody | Manufacturer (Catalogue number/ Reference) | Host | Working dilution | | |
|---|---|---|---|---|---|
| | | | IF | EM | IB |
| **alpha actin** | Sigma-Aldrich (A4700) | Mouse mAb | | | 1:500 |
| **GFP (B-2)** | Santa Cruz Biotechnologies (sc-9996) | Mouse mAb | 1:200 | | 1:200 |
| **LaminA/C** | Santa Cruz (sc-20681) | Rabbit pAb | | | 1:2,000 |
| **LaminB1** | Sigma-Aldrich (ZRB1143) | Rabbit mAb | 1:100 | | 1:5,000 |
| **LaminB3** | Own synthesis (Schütz et al., 2005) | Rabbit pAb | 1:100 | | |
| **13d4 (LAP2)** | Own synthesis (Alsheimer et al., 1998) | Mouse mAb | Undiluted hybridoma cell supernatant | | |
| **Myc** | Invitrogen (9E10) | Mouse mAb | 1:200 | | 1:8,000 |
| **PDI RL90** | Invitrogen (MA3-019) | Mouse mAb | 1:200 | | 1:1,000 |
| **Sun1** | Own synthesis (Göb et al., 2010) | Guinea pig pAb | 1:800 | | 1:5,000 |
| **Sun3** | Own synthesis (Göb et al., 2010) | Guinea pig pAb | 1:1000 | | 1:5,000 |
| **Sun4 peptide** | Own synthesis (Pasch et al., 2015) | Rabbit pAb | | | 1:5,000 |
| **Sun4-NTD** | Own synthesis (Pasch et al.,2015) | Guinea pig pAb | 1:2000 | 1:400 | 1:5,000 |
| **Sun4-NTD** | Own synthesis (Pasch et al.,2015) | Rabbit pAb | | 1:2400 | |
| **IgG** | Jackson ImmunoResearch Laboratories (011-000-003) | Rabbit pAb | | | |
| **IgG** | Jackson ImmunoResearch Laboratories (006-000-003) | Guinea pig pAb | | | |

**Table S4: Secondary antibodies used in this study.**

| Antibody | Manufacturer (catalogue numer) | Host | Antigen | Dilution |
|---|---|---|---|---|
| **6nm Gold** | Jackson ImmunoResearch Laboratories (706-195-148) | Donkey | Guinea pig IgG | 1:10 |
| **6nm Gold** | Jackson ImmunoResearch Laboratories (111-195-144) | Goat | Rabbit IgG | 1:10 |
| **HRP** | Jackson ImmunoResearch Laboratories (106-035-003) | Goat | Guinea pig IgG | 1:10,000 |
| **HRP** | Jackson ImmunoResearch Laboratories (706-035-148) | Donkey | Guinea pig IgG Cross-adsorbed | 1:10,000 |
| **HRP** | Jackson ImmunoResearch Laboratories (111-035-003) | Goat | Rabbit IgG | 1:10,000 |
| **HRP** | Jackson ImmunoResearch Laboratories (711-035-152) | Donkey | Rabbit IgG Cross-adsorbed | 1:25,000 |
| **HRP** | Jackson ImmunoResearch Laboratories (115-035-003) | Goat | Mouse IgG | 1:10,000 |
| **Alexa 488** | Invitrogen (A-11008) | Goat | Rabbit IgG | 1:200 |
| **Alexa 488** | Invitrogen (A-10680) | Goat | Mouse IgG | 1:300 |
| **Texas Red** | Jackson ImmunoResearch Laboratories (115-075-044) | Goat | Mouse IgG | 1:150 |
| **Texas Red** | Jackson ImmunoResearch Laboratories (106-075-003) | Goat | Guinea pig IgG | 1:50 |
